# Characterization of a novel *Lbx1* mouse loss of function strain

**DOI:** 10.1101/2021.08.25.457618

**Authors:** Lyvianne Decourtye, Jeremy A. McCallum-Loudeac, Sylvia Zellhuber-McMillan, Emma Young, Kathleen J. Sircombe, Megan J. Wilson

## Abstract

Adolescent Idiopathic Scoliosis (AIS) is the most common type of spine deformity affecting 2-3% of the population worldwide. The etiology of this disease is still poorly understood. Several GWAS studies have identified single nucleotide polymorphisms (SNPs) located near the gene *LBX1* that is significantly correlated with AIS risk. LBX1 is a transcription factor with roles in myocyte precursor migration, cardiac neural crest specification, and neuronal fate determination in the neural tube. Here, we further investigated the role of LBX1 in the developing spinal cord of mouse embryos using a CRISPR-generated mouse model expressing a truncated version of LBX1 (*Lbx1*^*Δ*^). Homozygous mice died at birth, likely due to cardiac abnormalities. To further study the neural tube phenotype, we used RNA-sequencing to identify 410 genes differentially expressed between the neural tubes of E12.5 wildtype and *Lbx1*^*Δ/Δ*^ embryos. Genes with increased expression in the deletion line were involved in neurogenesis and those with broad roles in embryonic development. Many of these genes have also been associated with scoliotic phenotypes. In comparison, genes with decreased expression were primarily involved in skeletal development. Subsequent skeletal and immunohistochemistry analysis further confirmed these results. This study aids in understanding the significance of links between Lbx1 function and AIS susceptibility.

## Introduction

Adolescent Idiopathic Scoliosis (AIS) is a 3-dimensional deformity of the spine and trunk and affects millions of people. It is characterized by a lateral curvature of the spine with a Cobb angle greater than 10° and does not result from either injury or vertebral malformation (Weinstein et al., 2008). This coincides with a rotational deformity of the vertebral column, even though the vertebrae and associated paraspinal musculature appear normal. AIS emerges between 11 and 18 years old in healthy children with no previous structural problems of the spine (Fadzan and Bettany-Saltikov, 2017). Scoliosis is often debilitating, with patients developing difficulties with movement and breathing. However, the etiology of this disease remains poorly understood. Various theories involve a genetic, neuromuscular, or biomechanical disorder. AIS has notably be considered as an abnormal growth of the skeleton (Yim et al., 2012), a ciliary defect (Grimes et al., 2016), and more recently as an alteration of the proprioceptive system (Assaiante et al., 2012; Blecher et al., 2017).

Due to the inheritability of the disease, genome-wide gene association studies (GWAS) have been performed. These studies have identified genes associated with AIS that function in processes such as cell adhesion and axon guidance (Fadzan and Bettany-Saltikov, 2017). These studies have notably identified three single nucleotide polymorphism (SNP) regions on chromosome 10q24.31 located within non-coding sequences downstream of the Ladybird homeobox 1 (*LBX1*) gene, an evolutionally conserved transcription factor (Grauers et al., 2015; Liu et al., 2017; Londono et al., 2014; Takahashi et al., 2011). One such SNP is r1190870, located in a region that physically interacts with the *Lbx1* promoter region. *In vitro* reporter studies predicted that the risk SNP (T) increases transcriptional activity (Guo et al., 2016). Thus, it has been suggested that the risk SNP confers AIS susceptibility through increased expression of *Lbx1*. Moreover, the ubiquitous overexpression of human LBX1 induced body curvature in zebrafish embryos (Guo et al., 2016). This previous research has led to LBX1 as a top candidate to functionally study concerning understanding AIS disease etiology.

LBX1 is essential for normal embryonic development and is required for the correct migration of muscle precursor cells and acquisition of the dorsal identity of forelimb muscle, specification of cardiac neural crest, and neuronal identity determination in the dorsal horn of the developing neural tube (Kruger et al., 2002; Martin and Harland, 2006; Masselink et al., 2017; Schafer et al., 2003). In addition, in the spinal cord, LBX1 is essential for the specification of GABAergic interneurons and the development of appropriate somatosensory pathways, leading to the correct distribution of sensory and motor information paraskeletal muscle (Cheng et al., 2005; Sieber et al., 2007).

The LBX1 protein is characterized by an engrailed homology domain (EH1) and a homeobox DNA-binding domain (Martin and Harland, 2006; Mizuhara et al., 2005). The EH domain binds transducing-like enhancer of split (Gro/TLE) proteins in mammals. Studies in mice revealed TLE genes are expressed from E7-8 in the neural tube, and expression continues throughout development in the CNS, axial skeleton, musculature, and neural crest-derived tissues (Dehni et al., 1995; Koop et al., 1996). *In vitro* studies have shown that LBX1 binds with Gro/TLE factor (Grg1) via the LBX1 EH1 domain to repress transcription. Loss of the EH1 domain prevents this protein-protein interaction and relieves repression (Mizuhara et al., 2005). Groucho-related genes (Grg) are expressed in the ventral region of the E10.5 neural tube (Muhr et al., 2001). In addition to TLE proteins, SKI family transcriptional co-repressor 1 (SKOR1), also known as Co-repressor to LBX1 (Corl1), binds to LBX1 but independently of the EH1 domain. SKOR1 is co-expressed in dorsal interneurons (dl4 and dl5) and late born dlLA and dlLB interneurons with LBX1 (Mizuhara et al., 2005). Together, this suggests two different mechanisms through which LBX1 represses gene expression, one of which is dependent upon the EH1 domain.

Our study aims to extend our knowledge on *Lbx1* by using a novel mouse model generated by CRISPR-CAS9 gene editing: *Lbx1*^*Δ*^. This is predicted to create a truncated version of the LBX1 protein, lacking the EH1 domain to alter expression of Lbx1 target genes. First, RNA-sequencing analysis was performed on E12.5 neural tubes for WT and *Lbx1*^*Δ*^ littermates, followed by gene ontology analysis. Subsequently, phenotyping of the E18.5 embryo was undertaken to examine the downstream consequences of *Lbx1*^*Δ*^ on embryonic development.

## Materials & Methods

### Animal information

To study Lbx1 function, we generated an *Lbx1* edited line named *Lbx1*^*Δ*^ using the CRISPR-Cas9 gene-editing system. This mouse line was developed at the Australian Biomedical Resources facility (Sydney, Australia) and had a deletion of 7 bp (AAGCCGC) in the coding sequence of *Lbx1*, inducing a stop codon, resulting in the deletion of the N-terminus portion of the LBX1 protein (Fig. S1; Fig. 1A). Mice were housed under standard conditions in individually ventilated cages. Heterozygous *Lbx1*^*Δ*^ (*Lbx1*^*Δ/+*^) mice were mated, the presence of a copulation plug was assessed at the commencement of the light cycle, and its appearance was considered as 0.5 days *post-coitus* (dpc). Females were then euthanized by cervical dislocation at embryonic development E12.5, E15.5, and E18.5 depending on the experiment. Embryos were then dissected, and tissue harvested and was stored either fresh frozen or, if used for future RNA extraction, in RNAlater (ThermoFisher). All animal procedures were performed with the approval of the University of Otago Animal Ethics Committee.

**Figure 1.**
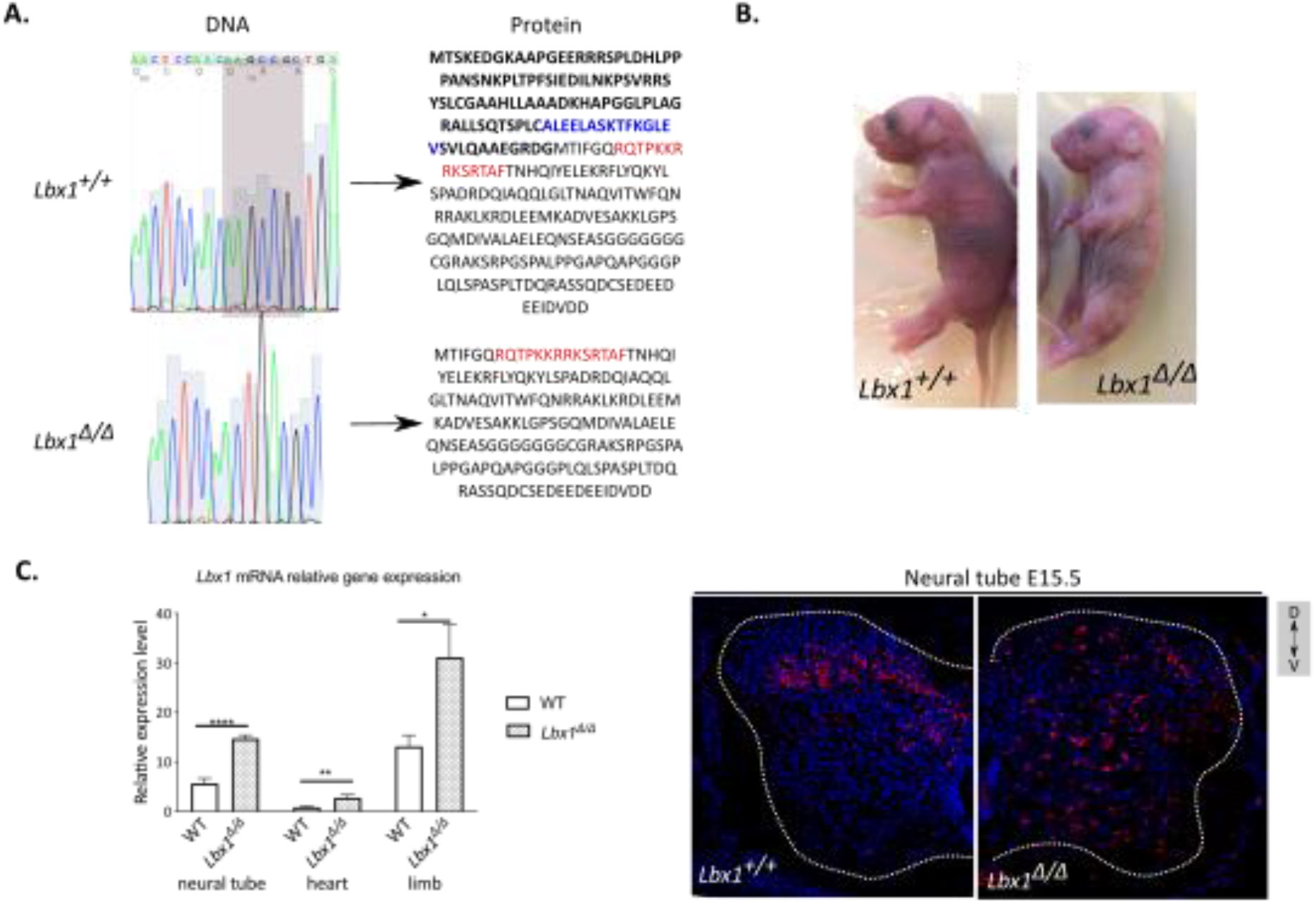
Impact of the CRISPR-cas9 deletion on *Lbx1* expression. (**A**) Comparison between Lbx1 coding DNA sequence of a WT embryo (control) and an *Lbx1*^*Δ/Δ*^ mutant. The gene edit resulted in a 7 bp deletion and is predicted to produce a truncated LBX1 protein without the N-terminal part of the protein (in bold). Predicted NLS is shown in red, NES is shown in blue. (**B**) *Lbx1* mRNA expression in WT and *Lbx1*^*Δ/Δ*^ E15.5 embryos in the neural tube, heart, and limb (n=6-9 per group). (**C**) Immunofluorescence of LBX1 protein (in red) and nuclei labeling with DAPI (in blue) in WT and *Lbx1*^*Δ/Δ*^ E15.5 neural tubes. V: ventral, D: dorsal.

### Genotyping

Mice ear notches or embryo tail tips were genotyped by PCR using Kapa2G Robust HotStart PCR Kit (Kapa Biosystems, KK5517) and *Lbx1* primers (Table S1). Excess of primers and dNTPs were then removed using ExoSAP-IT PCR Cleanup Reagents (ThermoFisher). Samples (3 ng) were finally sequenced on a 3730xl DNA Analyser (ThermoFisher) by the Genetic Analysis Services of the Otago University. Sequences were viewed using the 4Peaks application v1.7.1. Mice and embryos were subsequently considered as control (WT), heterozygote (*Lbx1*^*Δ/+*^), or homozygotes (*Lbx1*^*Δ/Δ*^).

### Whole Mount Skeletal Staining

Skeletal staining of E18.5 embryos was performed as previously described (Catela et al., 2009; Rigueur and Lyons, 2014). In brief, embryos had their skin removed, eviscerated, and fixed in 95% ethanol (EtOH) overnight (O/N). Embryos were then placed in acetone overnight to permeabilize cells and dissolve fat and subsequently transferred to alcian blue for cartilage staining O/N (0.03% alcian blue, 80% EtOH, 20% glacial acetic acid). Samples were then twice washed with 70% EtOH and finally in 95% EtOH O/N. Embryos were then cleared for 1.5 h in 1% KOH and left for 4 h in alizarin red for bone staining (0.005% alizarin red, 1% KOH). Clearing of the embryos was completed in KOH 1%/Glycerol 50% O/N and embryos were finally stored in glycerol. Digital images were obtained using a Nikon camera and a macro lens, and higher magnification images were performed using Leica microscope M80 studio and a MC170 HD camera. Final measurements were done using ImageJ (v1.52k).

### Histological analysis

For paraffin embedding and histology, E15.5 embryos were fixed overnight in 4% PFA. The skin was removed from E18.5 embryos, and the forelimb and head were removed before fixation in formalin for 3 days. The embryos were then washed in 70% EtOH before embedding in paraffin and sliced at 5 µm. Tissue morphology of E15.5 and E18.5 embryos were observed using hematoxylin and eosin (H&E) staining; embryos were sectioned at 5µm thickness. After deparaffinization and rehydration, slides were stained with hematoxylin for 4 min followed by eosin for 30 s.

For Masson trichrome (MT3) staining, after deparaffinization and rehydration, sections were stained in celestine blue (5 min), followed by hematoxylin (4 min), 0.5% acid fuchsin (5 min), 1% phosphomolybdic acid (4 min), 2% methyl blue (3.5 min) and finally, 1% acetic acid (2 min). These three dyes selectively stain muscle (red), collagen (blue), and nuclei (black).

Cresyl violet and Nissl stains were used to identify the neuronal structure in the neural tube. Cresyl violet stains the Nissl substance in the cytoplasm of neurons, a granular purple-blue while the background stays unstained. Briefly, after deparaffinization and rehydration, slides were stained in cresyl violet for 20 min, followed by 0.25% glacial acetic acid. At the end of the staining steps, slides were mounted in DPX and visualized with an Olympus BX61 Montaging Light Microscope. WT and *Lbx1*^*Δ*^ embryos were then compared at the same level across the rostrocaudal axis.

The EHistology Atlas was used as a resource for embryos analysis (http://www.emouseatlas.org/emap/eHistology/) (Richardson et al., 2014). ImageJ with Cell Counter plugin was used to assess the number of neurons (stained with cresyl violet) in the dorsal part of the neural tube in controls and *Lbx1*^*Δ/Δ*^ embryos. For each individual, the number of cells was assessed in a specific size area on 4 different slides separated by 5um of thickness, and the average was taken to avoid any bias. Results were compared by genotype (n = 4-6 per group).

### Immunofluorescence

For Immunofluorescence, E15.5 embryos were fixed 3 hours in 4% paraformaldehyde (PFA), followed by 2 washes in phosphate-buffered saline (PBS, 1X). Embryos were then transferred, subsequently, in sucrose 15%:PBS and sucrose 30%:PBS at 4°C until the embryo sank. Samples were then placed in a 1:1 mixture of OCT and sucrose 30% for 30 min and embedded in OCT. Cryosections of 15 μm were used for experiments.

For immunofluorescence, slides were fixed in 4% PFA for 30 min and then incubated with a guinea pig anti-Lbx1 (gift from Pr. Carmen Birchmeier and Dr. Thomas Müller) diluted at 1/20,000, O/N at 4°C. Slides were then incubated with a Goat anti-Guinea pig IgG (Alexa fluor 568, Abcam, #ab175714), mounted with fluoroshield mounting medium with DAPI (Abcam, #ab104139), and visualized with a Nikon A1+ inverted confocal laser scanning microscope and a Nikon DSiR2 color 16MP camera.

### Immunohistochemistry

Primary antibodies used were as follows: Peripherin (1 in 600; Abcam #ab246502), Pax3 (1 in 10; DSHB #Pax3-s),NF-H (1 in 100; DSHB #R797.s), WT1 (1 in 300; Abcam #ab89901) and ISL-1/2 (1 in 5; DSHB #39.4). Secondary antibodies used were: goat anti-mouse horse radish peroxidase (HRP) (1 in 600; Abcam #ab97023), goat anti-rabbit HRP (1 in 1000; #ab6721).

Slides were deparaffinized and rehydrated through a xylene/ethanol series. Antigen retrieval was performed with a sodium citrate buffer 30 min microwave (medium-power) and allowed to cool for 20 min before washing with PBSx (PBS plus 0.025% TritonX-100), followed by blocking for 1 h (blocking buffer PBS, 0.1% Triton X100, 10% Goat serum, 0.1% BSA). The primary antibody was diluted in the blocking buffer and incubated overnight at 4°C. Following washing with PBSx, slides were incubated with secondary antibody for 2 h at room temperature before DAB reaction using the DAB detection kit (Abcam #ab64238). Slides were counterstained with hematoxylin to stain nuclei. IHC experiments were repeated on at least 3 biological replicates to confirm the expression patterns were reproducible.

### RNA Sequencing analysis

Total RNA of E12.5 neural tube tissue was extracted using the Ambion Purelink RNA mini kit according to the manufacturer’s instructions (ThermoFisher). Invitrogen Purelink DNase (ThermoFisher) was used to remove any remaining genomic DNA. RNA concentration and quality were checked on a nanodrop (average concentration obtained 700 ng/µl, ratio 260/280 > 2, ratio 260/230 > 1.9). The RIN number was then assessed with a Bioanalyzer (RIN obtained >9.2). Control or *Lbx1*^*Δ/Δ*^ neural tubes from the same litter were pooled together and were considered as one replicate. The library preparation and RNA-sequencing were performed on an Illumina HiSeq platform generating 150 bp paired-end reads (NEBNext Ultra RNA Library prep kit for Illumina). Two replicates per group were analyzed.

Initial analysis was performed by using Galaxy (https://usegalaxy.org.au). The bioinformatic pipeline is shown in Figure S2. Quality control of the reads (around 44 million reads per replicate, fastq files) was assessed with FastQC (v. 0.72). Sequence data were then trimmed and filtered using FastQ quality trimmer (v. 1.0.0) and Filter FastQ (v. 1.1.1), respectively. The pre-processed reads (fastq files) were then mapped to the mouse reference genome mm9 (UCSC) using Tophat (v. 2.1.1) producing bam files (mapping statistics in Table S2). The alignment of the reads was then checked on UCSC using BamCoverage (v. 3.1.2.0.0) to create bigwig files. Finally, featureCounts (v. 1.6.0.1) was used to assign reads to genes, and the results were combined using Generate count matrix (v. 1.0).

Differential expression analysis was performed on RStudio using EdgeR (v. 3.4.3; (Robinson et al., 2010)). Differentially expressed genes (DEGs) were considered significant when the False Discovery Rate (FDR) adjusted p-value was <0.05. The heatmap was generated using heatmap.2{gplots} (Warnes et al., 2009).

Gene ontology (GO) was assessed using gProfiler using default settings (Raudvere et al., 2019). Raw and processed data have been deposited in the NCBI gene expression omnibus (GEO) accession number GSE132271.

### Measurement of gene expression

Total RNA was extracted from neural tube, heart, and limb samples using Direct-zol RNA MiniPrep Plus (Zymo Research) according to the manufacturer’s instructions, and concentrations were assessed on nanodrop. A total of 1 µg of RNA was reverse transcribed using iScript reverse transcription supermix (Bio-Rad). Real-time PCR was performed as triplicates on 2 ng of cDNA using SYBR Green master mix (ThermoFisher) on a ViiA 7 real-time PCR System (ThermoFisher). The list and sequences of the primers used are presented in Table S1. The abundance of mRNA was measured using the comparative Ct method and normalized against *Gapdh* and *Actinb* reference genes.

### Statistical analysis

All data are presented as the mean ± SEM. Analysis was performed using Prism 7 software. Comparisons between WT (control) and Lbx1Δ were determined using a non-parametric Mann-Whitney test. Statistical significances of the fold changes were ascertained using a student’s t-test. Correlation analysis was analyzed using Pearson correlation coefficients. Statistically significant results are labelled as: * p<0.05, ** p<0.01, *** p<0.001.

## Results

### CRISPR-Cas9 *Lbx1*^*Δ*^ generation

To further investigate the role of LBX1, we generated a CRISPR-Cas9 mouse line, named *Lbx1*^*Δ*^, with a deletion of 7 bp (AAGCCGC) in the coding sequence of the *Lbx1* gene creating an early stop codon (Fig. 1A and Fig. S1). This deletion is predicted to produce a transcript with two protein products – 30 amino acid (aa) and a truncated 170 aa protein LBX1 protein without the EH domain at its N-terminus and leaving the DNA binding domain intact (Fig. 1A). Despite the absence of the EH domain, the truncated LBX1Δ is still predicted to bind DNA but would not recruit Groucho/TLE protein partners. This CRISPR-Cas9 mouse model allows us to study downstream processes affected by the absence of LBX1 protein-protein interactions. Lbx1Δ is still predicted to bind some co-repressors such as CORL1 (Mizuhara et al., 2005).

Heterozygous *Lbx1*^*Δ*^ (*Lbx1*^*Δ/+*^) did not present any obvious phenotype and were viable. However, homozygous *Lbx1*^*Δ*^ (*Lbx1*^*Δ/Δ*^) died at birth. This has also been observed with previous *Lbx1* knockout mice models (Brohmann et al., 2000; Hernandez-Miranda et al., 2018; Pagliardini et al., 2008). Therefore, for the rest of the experiments, only WT and homozygous *Lbx1*^*Δ*^ are considered.

*Lbx1* mRNA was first assessed in E15.5 embryos by RT-qPCR in three tissues known to require LBX1 function: the neural tube, the heart, and the forelimb. *Lbx1*^*Δ/Δ*^ exhibits a significant increase in *Lbx1* mRNA expression in all three tissues compared to WT embryos (Fig. 1B). This increase in *Lbx1* transcript levels has been observed in another *Lbx1* knockout model, suggesting a feedback loop with LBX1 controlling its own expression levels (Schafer et al., 2003).

To study the localization of LBX1 protein in the developing neural tube, immunofluorescence was performed on E12.5 and E15.5 cryosections, with a polyclonal antibody raised against the full-length protein. In WT embryos, LBX1 is strongly detected in the dorsal part of the neural tube (as previously described (Muller et al., 2002) (Fig. 1D). Furthermore, LBX1 expression in the neural tube is nuclear, co-localizing with the DAPI (Fig. 1D). In *Lbx1*^*Δ/Δ*^ E15.5 embryos, the staining for LBX1 protein is considerably weaker in the neural tube but still detectable and appears more widely distributed (Fig. 1E). This indicates that a protein is being produced in the *Lbx1*^*Δ/Δ*^ embryo.

### *Lbx1*^*Δ/Δ*^ mutation is embryonic lethal

To assess the impact of the LBX1^Δ^ truncation, we performed H&E and Trichrome (MT3) staining on E15.5 and E18.5 embryos (Fig. 2). H&E staining was used to assess the general morphology of tissue, whereas MT3 was used to selectively demonstrate muscle, collagen fibers, fibrin, erythrocytes, and nuclei. We examined more closely the morphology of the neural tube, the heart, and the forelimb, which is sensitive to the loss of LBX1 function (Brohmann et al., 2000; Muller et al., 2002; Schafer et al., 2003). Histology on E15.5 and E18.5 embryos showed that the overall morphology of the spinal cord seems to be similar between control and *Lbx1*^*Δ/Δ*^ (Fig. 2A and 2D; 2G and 2J). The grey matter is well defined in both genotypes and the subarachnoid spaces are consistent. At this age, *Lbx1*^*Δ/Δ*^ embryos appear to have abnormalities in the heart development with thickened left ventricular walls and an enlargement of the atria with abnormalities of the semilunar valves (Fig. 2B and 2D; 2H and 2K). Analysis of E15.5 and E18.5 forelimbs showed that in WT embryos, muscle groups are well defined with clear borders between muscle groups; in *Lbx1*^*Δ/Δ*^ forelimb sections, distinct muscle groups were present at E18.5 (Fig. 2C and 2E; 2I and 2L). This suggests that the N-terminal portion of LBX1 is dispensable for myoblast migration into the developing limb.

**Figure 2.**
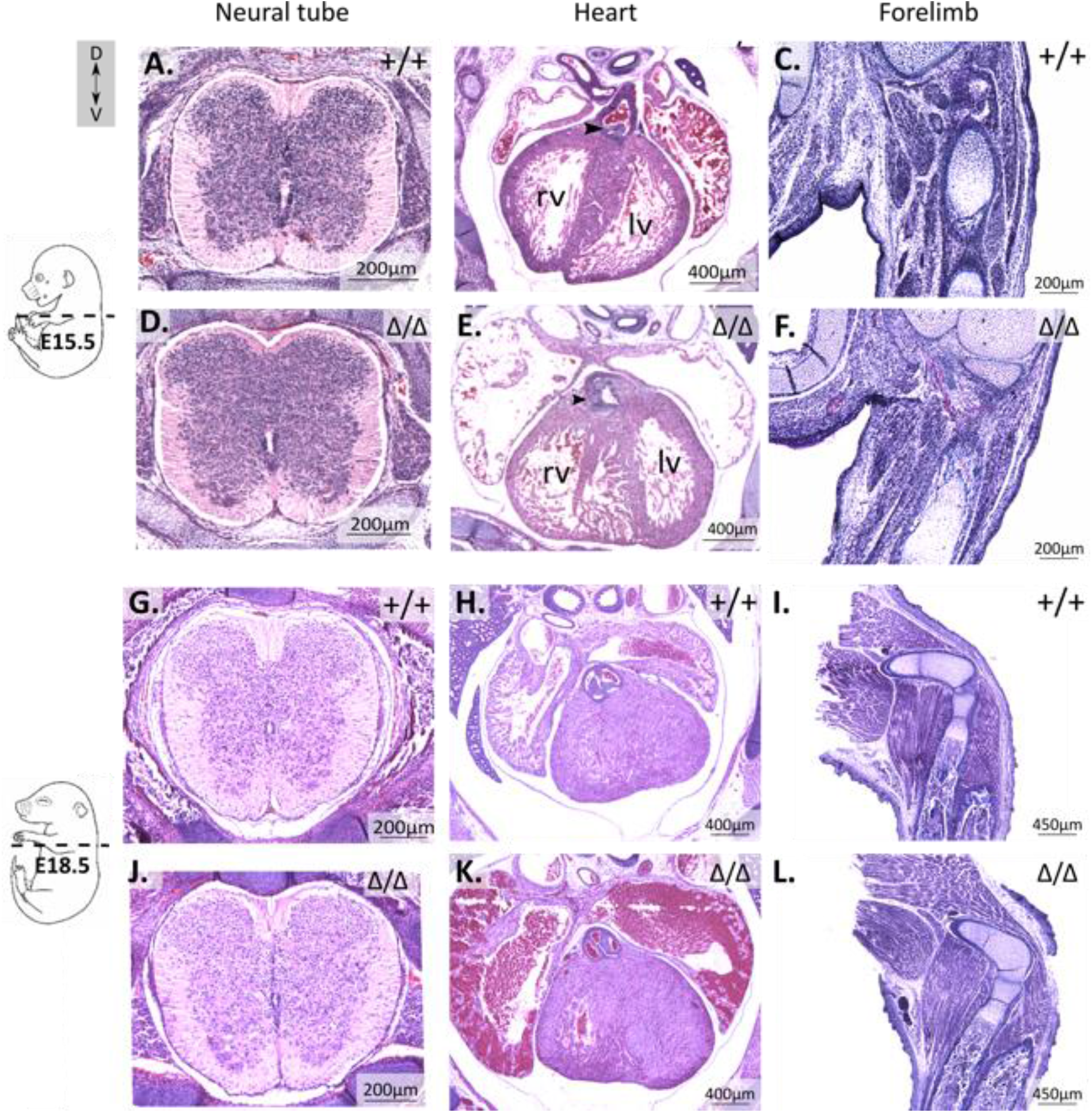
*Lbx1*^*Δ/Δ*^ interferes with embryonic development. Comparison of different tissue morphology of E15.5 and E18.5 embryos. H&E staining was performed on E15.5 WT (+/+) (**A-B**) and *Lbx1*^*Δ/Δ*^ embryos (**D-E**). Forelimbs were stained with Masson trichrome, in WT (**C**) and *Lbx1*^*Δ/Δ*^ (**F**). Neural tube morphology at E15.5 of controls (**A**) and *Lbx1*^*Δ/Δ*^ (**D**) looks similar. Hearts of *Lbx1*^*Δ/Δ*^ (**E**) have thicker ventricular walls and abnormalities of the semilunar valves (arrowheads) compared to controls E15.5 embryos (**B**). Forelimbs of E15.5 (**C** and **F**) and E18.5 (**F** and **L)** for both WT (**C** and **F**) and *Lbx1*^*Δ/Δ*^ (**I** and **L**) contain well-defined muscular groups. Abbreviations: rv: right ventricle, lv: left ventricle.

Earlier RT-qPCR analysis found *Lbx1* mRNA was overexpressed in the *Lbx1*^*Δ/Δ*^ heart and limb tissues at E15.5, a timepoint when *Lbx1* expression is expected to have reduced as it is several days since the completion of neural crest/myoblast migration and differentiation into respective structures. Heart tissue isolated from both WT and *Lbx1*^*Δ/Δ*^ embryos also showed significant increases in *MyoD* (7-fold, P=0.004), *Fhl2* (2-fold; P=0.0007), and *Pax3* (3-fold; P=0.01) mRNA expression (Fig. S3). Despite the overall gross normal appearance of muscle groups in the *Lbx1*^*Δ/Δ*^ limb, *MyoD* mRNA expression was significantly decreased ∼2-fold (P=0.03; Fig. S3), with no change to *Pax3* expression, or *Map2k1* (Fig. S3; P=0.88, P=0.15). This suggests that Lbx1Δ negatively impacts downstream processes in these tissues.

### RNA Sequencing analysis

To better understand the early deficit of *Lbx1* expression in the developing spinal cord, we performed an RNA-Sequencing (RNA-seq) analysis on E12.5 neural tubes (plus somite/neural crest) of *Lbx1*^*Δ/Δ*^ and WT littermates. At this age, *Lbx1* expression is easily detectible (data not shown) (Hernandez-Miranda et al., 2018). Two replicates per group were analyzed following the bioinformatic pipeline presented in Fig. S1. Following sequencing, the reads were mapped to the genome (UCSC_mm9) using Tophat, and differential expression analysis was performed using EdgeR. We found 410 genes were differentially expressed between WT and *Lbx1*^*Δ/Δ*^ embryos (adjusted p-value < 0.05), with 180 transcripts with higher expression and 230 transcripts with decreased expression in *Lbx1*^*Δ/Δ*^ neural tubes (Fig. 3A and 3B) (Table S3).

**Figure 3.**
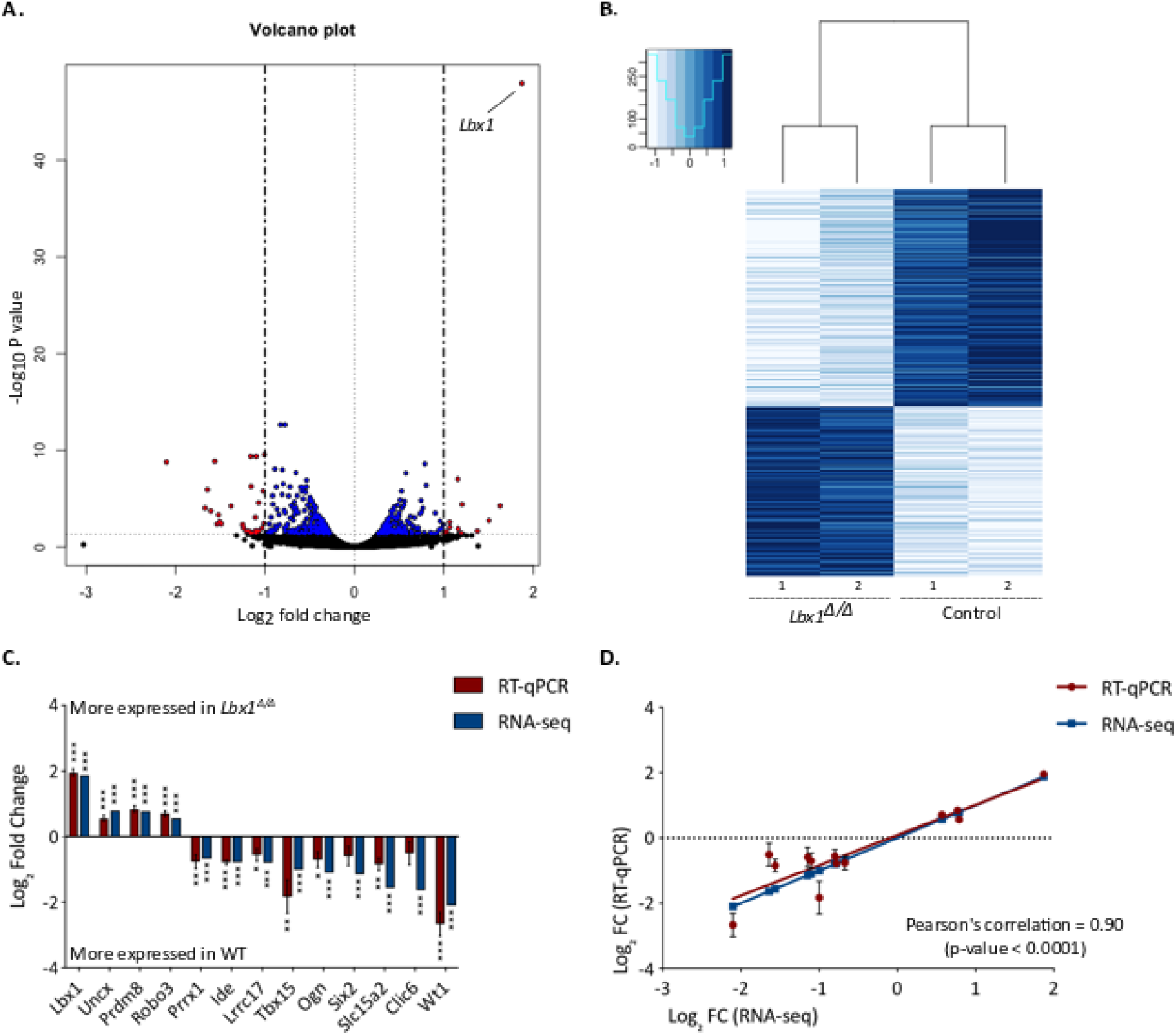
RNA-sequencing analysis of E12.5 neural tubes. (**A**) Volcano plot of the RNA-seq results between WT and *Lbx1*^*Δ/Δ*^ E12.5 neural tubes with colour dots (blue and red) indicating genes with an adjusted p-value <0.05 and in red the differentially expressed genes with a log_2_ fold change <-1 or >1 (list of genes in Table S3). (**B**) Heatmap of the 410 genes differentially expressed with individual hierarchical clustering of controls and *Lbx1*^*Δ/Δ*^. The genes with the highest expression are shown in dark blue and the low expression in pale blue. (**C**) Bar plot presenting Log_2_ fold change ± SEM between RNA-seq results (in blue) and RT-qPCR ones (in red, n=9-12 per group). One sample t and Wilcoxon test P < 0.05 (*), P < 0.001 (**), P < 0.0001 (***). (**D**) Scatter plot of the log^2^ fold change in both analyses with a Pearson’s correlation = 0.90, P < 0.0001. The solid lane on this plot represents the linear regression of this correlation.

To confirm the results obtained, RT-qPCR analysis was performed on 13 genes found to be differentially expressed by EdgeR analysis on RNA collected from independent E12.5 neural tubes (n=9-12). All genes showed a very similar change in expression to that of the RNA-seq data (Fig. 3C). The results obtained with both methods are highly correlated, as shown on the scatter plot of the log_2_ fold change (Pearson’s correlation = 0.90, adjusted p-value < 0.0001 (Fig. 3D)). These results validated those obtained by the bioinformatic analysis.

The gene with the largest increase in expression in *Lbx1*^*Δ/Δ*^ samples was *Lbx1* (1.87 log_2_FC, adjusted P = 1.38×10^−48^). The gene with the largest reduction in mRNA levels was Wilms tumor 1 (*Wt1*) (−2.1 logFC; adjusted P (Padj) = 1.68×10^−09^), a key component of interneurons involved in locomotor circuitry (Haque et al., 2018; Schnerwitzki et al., 2018). In terms of the top significantly DEGs (Padj 1.13×10^−48^ - 8.56×10^−09^), we mostly find genes involved in neurogenesis such as *Lbx1*, UNC homeobox (*Uncx*, 0.79), which regulates the proliferation and survival of neurons, and Calpain 6 (*Capn6*, -0.83) that had been implicated in neurodegenerative processes such as Alzheimer’s, as well as skeletal muscle and heart development (Sammeta et al., 2010; Tonami et al., 2013) (Higuchi et al., 2005). We also found differential expression of genes involved in bone formation such as T-box 15 (*Tbx15*, -1.00), a gene that determines the number of mesenchymal precursor cells and chondrocytes in vertebral development, ALX homeobox 4 (*Alx4*, -1.16) known for its involvement in osteoblast regulation, and osteoglycin (*Ogn*, -1.10) required for osteoblastogenesis (Chen et al., 2017; Singh et al., 2005; Yagnik et al., 2012).

### Gene Ontology analysis of differentially expressed genes

To further characterize the differentially expressed genes (DEGs), gene ontology analysis of biological processes was performed on the 410 genes differentially expressed in *Lbx1*^*Δ/Δ*^ E12.5 neural tubes compared to the controls, using gProfiler (Raudvere et al., 2019).

Regarding the genes with increased expression, the top biological processes (BP) by significance were all associated with aspects of neurogenesis and neuronal development (Padj = 6.7×10^−18^) (Fig. 4A; File S2). The top cell component terms were linked to the axon and the synapse (Padj = 5.3×10^−6^ and 2.7×10^−5^). Pathway analysis revealed enrichment for Notch signaling pathway (Padj = 4.6×10-3) and axon guidance (Padj = 1.31×10-2) (Fig. 4A; File S2). Thirty-two genes were associated with abnormal nervous system physiology in humans (Padj = 1.97×10^−2^; Fig. 4A, File S2).

**Figure 4.**
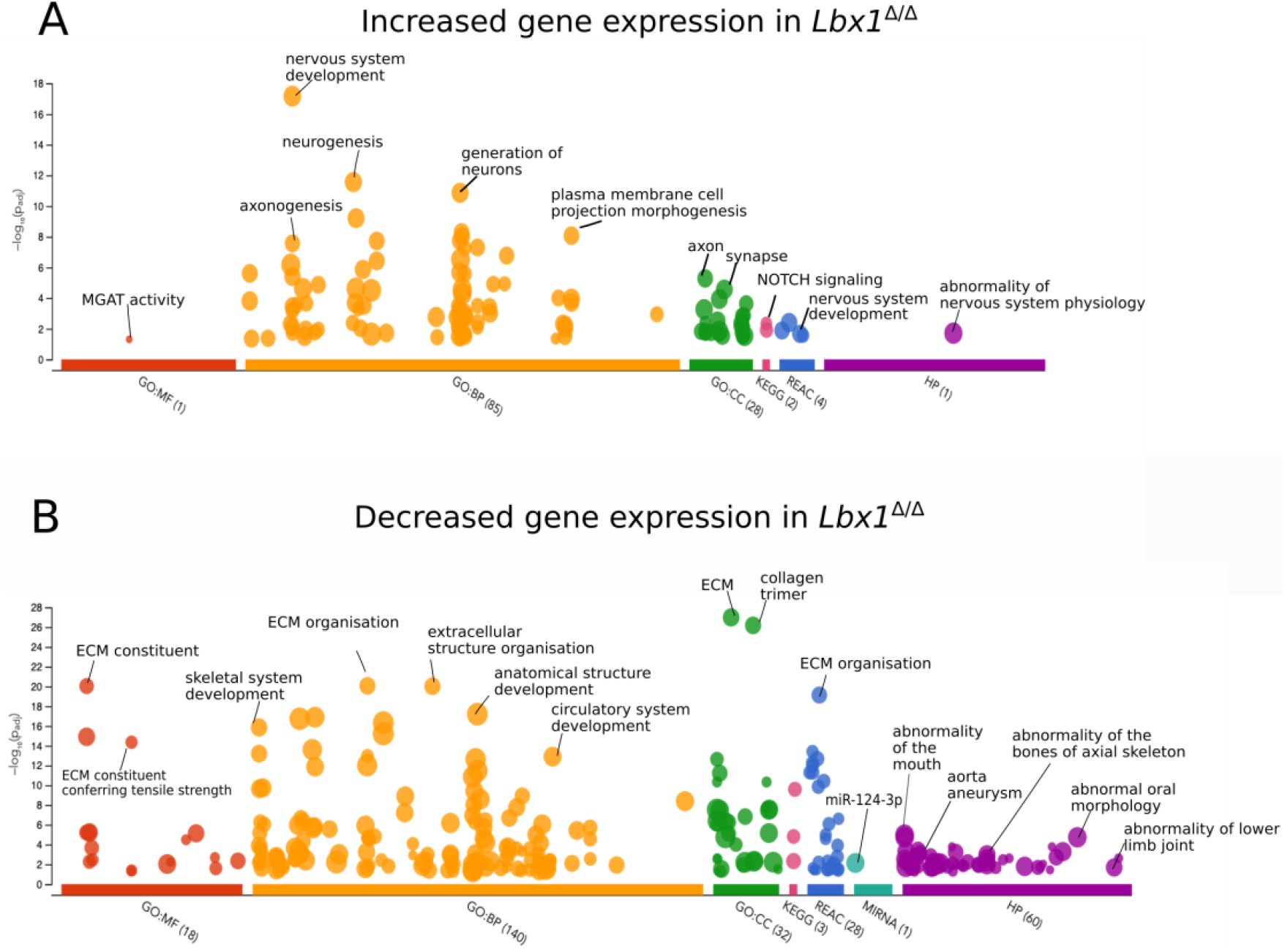
Gene ontology of differentially expressed genes. Manhattan plots separate the analysis into different categories: Gene ontology terms (GO) biological processes (BP), molecular functions (MP), cell component (CC), KEGG, REAC, miRNA targets, and Human phenotypes (HP). Adjusted p-values (Bonferroni correction) shown on the Y-axis by using gProfiler. (**A**) Profiling of genes with increased expression in *Lbx1*^*Δ/Δ*^ compared to WT. (**B**) Enrichment of gene ontology terms associated with downregulated genes in *Lbx1*^*Δ/Δ*^ neural tubes Abbreviations: MGAT (alpha-1,6-mannosylglycoprotein 6-beta-N-acetylglucosaminyltransferase), ECM (extracellular matrix).

Genes with decreased expression were significantly associated with the extracellular matrix (ECM); Padj = 8.77×10-^21^), skeletal development (Padj = 1.42×10^−16^) and circulatory system development (Padj = 1.27×10^−13^; Fig. 4B). Significant human phenotypes associated with these genes included abnormalities of mouth morphology, heart defects such as mitral valve prolapse (Padj = 0.0006), and skeletal issues (eg aplasia/hypoplasia affecting bones of the axial skeleton (Padj = 0.001) (Fig. 4B; File S2).

Additionally, within the differentially expressed genes in our mouse line, we found 21 genes that have previously been linked to scoliotic phenotypes (Table S5), such as *Notch1* (0.38, adjusted P < 0.001) and *Gpr26* (1.19, adjusted P < 0.05). Loss of *NOTCH1* function induces contiguous vertebral segmentation, short stature, and scoliosis (Loomes et al., 2007). GPR26 mutation results in altered cartilage biology and spinal column development producing a scoliosis phenotype (Karner et al., 2015; Xu et al., 2015). Several DEGs are also associated with syndromic congenital scoliosis: *Dll3, Meox1, Myot, Sall1, Lfng* (Takeda et al., 2018) (Bulman et al., 2000) and Horizontal gaze palsy and progressive scoliosis (HGPPS): Robo3 (Jen et al., 2004; Volk et al., 2011) (Table S5). Many of these genes are also part of the Notch signaling pathway.

### Impaired neurogenesis in the *Lbx1*^*Δ*^ neural tube

These results show that an alteration of LBX1 function in the *Lbx1*^*Δ/Δ*^ E12.5 neural tube impacted embryonic and cellular biological processes, particularly neurogenesis and skeletal development. As these two processes have previously been linked to AIS (Blecher et al., 2017; Yim et al., 2012) we examined these aspects of the phenotype further.

To study the impact of *Lbx1*^*Δ*^ deletion on neurogenesis, the general aspect of the neural tube was assessed by Nissl staining on E15.5 and E18.5 embryos and the number of neurons (stained purple-blue) in the dorsal part of the neural tube was counted. No significant differences were observed between *Lbx1*^*Δ/Δ*^ and WT E15.5 dorsal horn neuron number (Fig. 5A and 5C). However, by E18.5, there was a significant decrease in *Lbx1*^*Δ/Δ*^ dorsal horn neuronal number compared to WT (−41.3 ± 3.45, p < 0.0001 (n=4-5); Fig. 5B-C)

**Figure 5.**
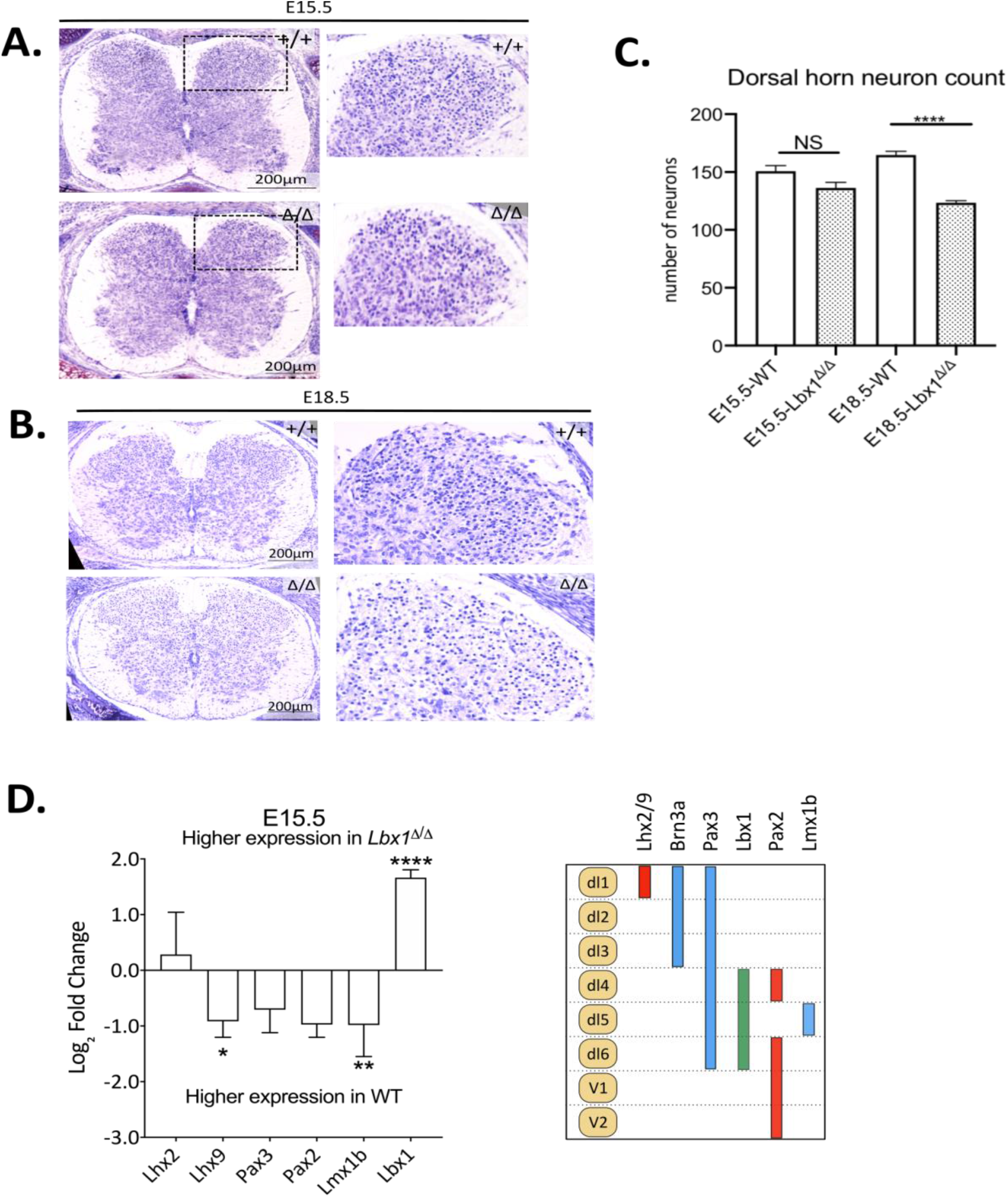
Neural tube neurogenesis in *Lbx1*^*Δ/Δ*^ embryos. (**A, B**) Nissl staining was performed on E15.5 and E18.5 embryos to assess the number of neurons in *Lbx1*^*+/+*^ and *Lbx1*^*Δ/Δ*^ E15.5 neural tubes (**A**) and *Lbx1*^*+/+*^ and *Lbx1*^*Δ/Δ*^ E18.5 neural tubes (**B**). The number of neurons was assessed within the dorsal horn. No significant differences in neuron numbers were found at E15.5, whereas at E18.5, neuron numbers were significantly decreased in *Lbx1*^*Δ/Δ*^ embryos. (**C**) The known pattern of expression of the different neuronal subpopulations according to the literature (Gross et al., 2002). (**D**) RT-qPCR analysis in E12.5 and E15.5 neural tubes. The expression of the neuronal subpopulation markers is globally increased at E12.5 and decreased at E15.5 in the mutant compared to the WT. The data shown are mean ± SEM. One sample t and Wilcoxon test P < 0.05 (*), P < 0.001 (**), P < 0.0001 (***).

To precisely study the different neuronal subpopulations in the *Lbx1*^*Δ/Δ*^ neural tube, qPCR and IHC were used. The neural tube is defined by neuronal layers (dl1 for the most dorsal layer to dl6 and V1, V2 for the ventral layers) (Fig. 5C). We used markers of the different neuronal layers to examine any changes in neuronal development of the neural tube. *Lhx2/9* is known to label the most dorsal interneurons population: dl1, while populations dl1-dl3 expresses *Brn3a, Pax3* dl1 - dl6, *Lbx1* dl4 - dl6, *Pax2* dl4 and dl6 - V2, and *Lmx1b* dl5 interneurons(Gross et al., 2002).

First, changes to mRNA expression levels were assessed by qPCR using E12.5 and E15.5 neural tubes from WT and *Lbx1*^*Δ/Δ*^ embryos. In E12.5 *Lbx1*^*Δ/Δ*^ embryos, we observed a general increase fold change of expression of the neuronal markers, with a significance for *Lhx2* (P = 0.03), *Lhx9* (P < 0.0001), *Brn3a* (P < 0.0001) and *Pax3* (P = 0.002) compared to the WT (Fig. 5D). Conversely, at E15.5, we observed decreased mRNA expression different markers, with a significant decrease for both *Lhx9* (P = 0.024) and *Pax2* (P = 0.006) (Fig. 5D).

Genes were arranged using the knowledge matrix of Delile et al., (2019), which defines genes known to contribute to each cell type (not all DEGs are in this matrix) (Delile et al., 2019) Fig. S4. Thus, DEGs are broadly expressed in multiple cell types, suggesting that *Lbxl*^*Δ*^ results in more general disruption of neurogenesis, not limited to the dorsal horn.

An essential function of interneurons is to coordinate incoming afferent signals and mediate their course peripherally. Peripherally projecting neurons can be stained for peripherin, a type III intermediate filament protein found in the distal regions of extending axons that mark neurons that project towards peripheral structures (Leonard et al., 1988; Troy et al., 1992) (Fig. 6A). In WT mice, we observed long axons extending through midline from the dorsal side of the spinal cord, where they make connections to their respective interneurons ((Lu et al., 2015); Fig. 6A asterisks). In comparison, *Lbx1*^*Δ/Δ*^ had darker stained shortened axons present through the midline region of the spinal cord (Fig. 6A, asterisks) with premature termination of axons. Peripherin actively stained the motoneuron population within the ventral horn, with obvious peripherally extending axons in the WT (6Ci, arrows). In the *Lbx1*^*Δ/Δ*^ section, there was a smaller motoneuron population and a distinct lack of these peripherally extending neurons expressing peripherin (Fig. 6Ai). Medial motor axons, projecting towards the ventral root and midline, were absent in the *Lbx1*^*Δ/Δ*^ spinal cord (Fig. 6Ai, arrows). In WT embryos, axons project from the dorsal root into the dorsal horn (Fig. 6Aii’ arrows); this is absent in the *Lbx1*^*Δ/Δ*^ dorsal horn. Overall, peripherin staining indicates a problem with axon outgrowth/extension for neurons with type III intermediate protein in the *Lbx1*^*Δ/Δ*^ mouse.

**Figure 6.**
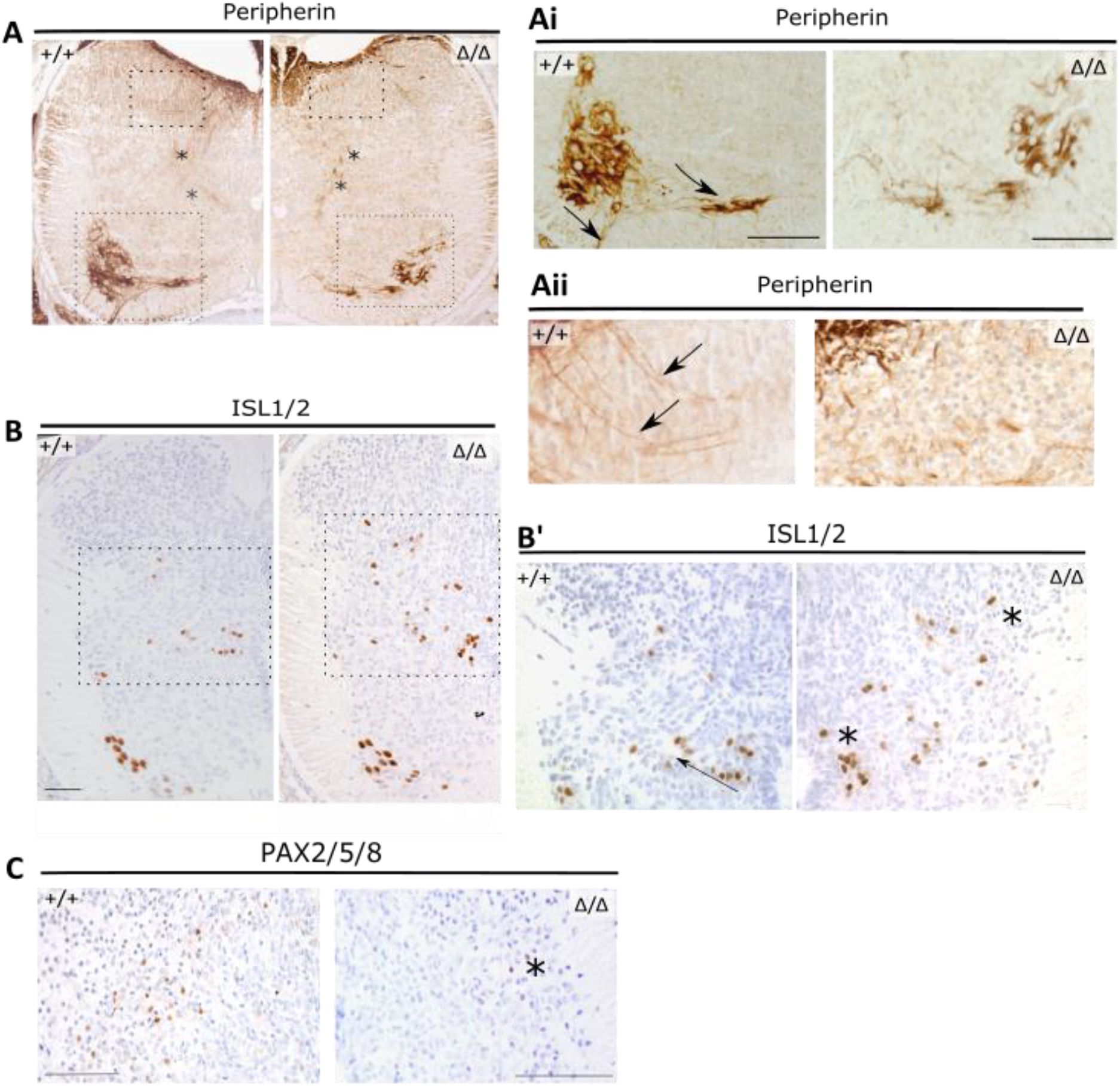
Later development of the spinal cord in WT and *Lbx1*^*Δ/Δ*^ embryos. E15.5 sections shown are from the thoracic region. **A**. Peripherin staining of WT (+/+) and *Lbx1*^*Δ/Δ*^ sections. Asterisks show axons through a central region of the spinal cord. Higher magnification image of the ventral horn (Ai). Arrows indicate axons projecting from the Peripherin positive motor neurons towards the midline and the ventral root. Higher magnification image of the dorsal horn of both WT and *Lbx1*^*Δ/****Δ***^ sections (Aii), arrows indicate axons projecting from the dorsal root into the dorsal horn of the spinal cord. **B**. IHC for ISL1/2 protein, which marks dl4, dl6 and ventral motoneurons. B’ is a higher magnification image of the region shown in B (dashed box). Asterisks indicate some abnormally migrating ISL1/2 neurons compared to the WT (arrow). **C**. PAX2/3/5 antibody detected expression within in the dl4 neurons, but not in the *Lbx1*^*Δ/Δ*^ embryo.

Sections from E15.5 embryos were also stained with ISL1/2 and PAX2/5 (Fig. 6B & C). Islet 1/2 (ISL1/2) is required for motor neuron development and is expressed in the ventral horn within motoneurons as well as the dl3 population (Fig.6B; (Gross et al., 2002; Pfaff et al., 1996)). We did not detect any difference in the number of ISL1/2 positive neurons (data not shown). However, they often migrated in a more random direction in the *Lbx1*^*Δ/Δ*^ spinal cord than WT sections (Fig. 6B’), suggesting a defect in axon guidance.

Staining for PAX2/3 protein in WT embryos was found towards the middle of the developing spinal cord, where dl4 and dl6 inhibitory interneurons express it (Fig. 6B, (Gross et al., 2002; Muller et al., 2002)). On the other hand, PAX2/5 protein expression was not detectable in E15.5 *Lbx1*^*Δ/Δ*^ embryo sections (Fig. 6B). This suggests a loss of dl5 and dl4 interneurons.

These results highlight changes to neurogenesis in *Lbx1*^*Δ/Δ*^ embryos and agree with the RNA-seq data, showing a modification of gene expression involved in both neuronal identity and connectivity and the phenotypic consequences of altered gene expression (Table S4).

### Skeletal malformation in *Lbx1*^*Δ/Δ*^ mice

Gene ontology analysis showed that in *Lbx1*^*Δ/Δ*^ neural tubes, 44.2% of the genes with decreased expression were linked to skeletal development (Fig. 7). To evaluate the impact on the developing skeleton, we performed whole-mount skeletal staining on E18.5 embryos with Alcian Blue to stain cartilage and alizarin red to stain for bones (Fig. 7). *Lbx1*^*Δ/Δ*^ embryos revealed hypoplasia of the skeleton with the overall length of the spine, measured from the first cervical vertebra to the end of the tail, was significantly smaller compared to WT littermates (Fig. 7A and 7E, 6.5% mean reduction in length; P = 0.04). Closer examination of the skull did not reveal any significant changes in the size, measured from either the nose to the end of the parietal bone (measure 1) or the end of the interparietal bone (measure 2) (Fig. 7B and 7F). At E18.5, the general size of the vertebrae was measured horizontally (measure 1) and vertically (measure 2) on the first lumbar vertebra (L1) and showed a significant reduction in size for *Lbx1*^*Δ/Δ*^ when compared to WT littermates (Fig. 7C and 7G, 11.7% (measure 1; P= 0.05) and 6.4% (measure 2; P= 0.03)). The length of the hindlimb was also measured, from the head to the end of the tibia (with the cartilage stained in blue, measure 1 and without the cartilage stained in red, measure 2). It showed a significant reduction in *Lbx1*^*Δ/Δ*^ embryos compared to WT for measure 1 (Fig. 7D and 7H; reduction of 10% (measure 1; P= 0.03) and 8.9 (measure 2; P = 0.11). These results are in agreement with the RNA-sequencing results and suggest that incorrect LBX1 function can change typical skeletal development.

**Figure 7.**
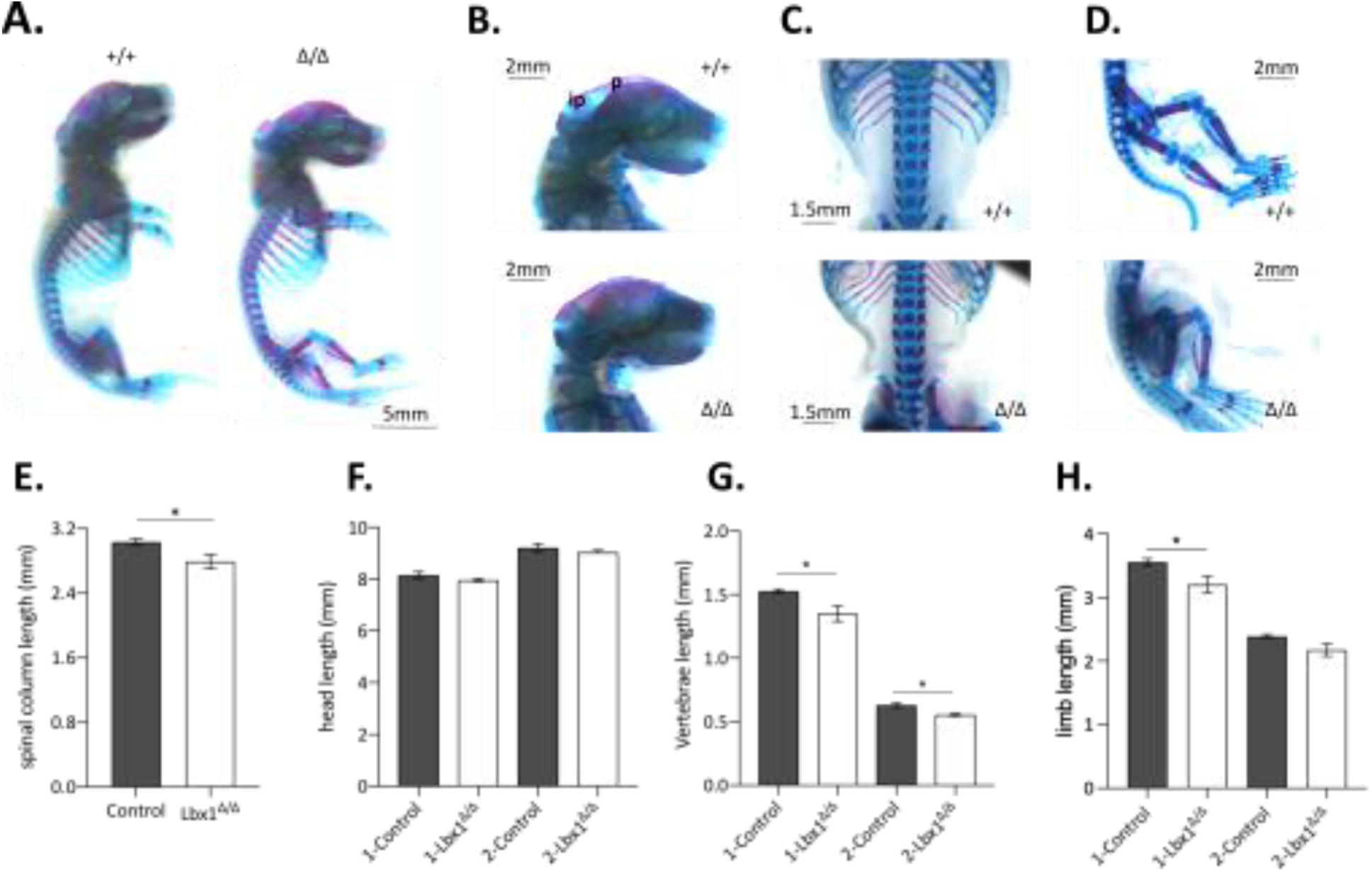
Skeletal anomalies in *Lbx1*^*Δ/Δ*^ embryos. (**A**) Comparison between whole-mount skeletal staining of WT (+/+) and *Lbx1*^*Δ/Δ*^ embryos (Δ/Δ) at E18.5. Cartilage is stained in blue and bones in red. (**B**) Lateral view of the skull with p: parietal and ip: interparietal bone. (**C**) Ventral view of the vertebrae and (**D**) lateral view of hindlimbs. (**E**) Bar plot representing the size difference of the spine between E18.5 WT and *Lbx1*^*Δ/Δ*^ embryos. (**F**) Head length analysis between the nose to the end of the parietal bone (1-Control and 1-*Lbx1*^*Δ/Δ*^) or the end of the interparietal bone (2-Control and 2-*Lbx1*^*Δ/Δ*^), no difference was observed. (**G**) Vertebrae analysis measured horizontally (1-Control and 1-*Lbx1*^*Δ/Δ*^) and vertically (2-Control and 2-*Lbx1*^*Δ/Δ*^) on the first lumbar vertebra (L1) showed significant hypoplasia in *Lbx1*^*Δ/Δ*^ compared to the controls. (**H**) The length of the hindlimb was finally measured, from the head of the tibia to its end (with the cartilage stained in blue: 1-Control and 1-*Lbx1*^*Δ/Δ*^ or without the cartilage: 2-Control and 2-*Lbx1*^*Δ/Δ*^) and showed a significant diminution in *Lbx1*^*Δ/Δ*^ embryos compared to WT. n=5 per genotype, mean with SEM is shown. The result of the Mann-Whitney two-tailed P-value is also shown when significant (* P < 0.05).

## Discussion

The present study aimed to investigate the role of LBX1 during embryonic development, to understand its association with AIS. The data presented in this study further supports the essential role of LBX1 in the developing spinal cord and that disruption to LBX1 functions alters the expression of a wide range of genes and biological processes that can be correlated with scoliotic phenotypes.

The truncated version of LBX1 lacks the EH1 domain needed for interaction with GRO/TLE factors (Dehni et al., 1995; Koop et al., 1996), although it may still bind to some co-repressors such as CORL1 (Mizuhara et al., 2005). Immunofluorescence showed that the LBX1Δ protein was still expressed and present in the nucleus. We also attempted a Western blot with embryo tissue, using both commercial antibodies and the antibody used herein IF. Unfortunately, none successfully produced a single on the blot for WT or truncated LBX1, indicating that these antibodies do not work well in a Western. RNA-seq analysis identified 410 differentially expressed genes in the deletion line compared to the control at E12.5. Gene ontology analysis revealed the biological processes altered in these mice included nervous system development (such as neurogenesis, synapse organization, and axon guidance) and ECM composition (File S2, Fig. 4).

The gene ontology analysis of the decreased gene expression found several downregulated biological processes in *Lbx1*^*Δ/Δ*^ tissues, including general tissue development, extracellular and cellular development, and embryonic skeletal system development along with ossification, bone morphogenesis, chondrocyte differentiation, and cartilage development. A more thorough analysis of the skeletal development showed that the general skeleton of *Lbx1*^*Δ/Δ*^ embryos is significantly smaller than the WT, including a decrease in the length of the spine, vertebrae, and the limb (Fig. 7). The alteration of the size of the limb could be explained by a reduction in prenatal movements, where appropriate nervous system development contributing to prenatal movements stimulates growth. Previously, immobilization due to reduced mechanical stimulation during prenatal development has been observed to result in short limbs (Nowlan et al., 2010; Pollard et al., 2017; Rodriguez et al., 1988; Rot-Nikcevic et al., 2006), a feature which is reversed following neuromuscular stimulation, resulting in increased in length in chick embryos (Heywood et al., 2005).

Lbx1-expressing cells migrate from the NC into the developing heart between E9-E10.5(Schafer et al., 2003). Although aberrant migration patterns were observed for Lbx1^+^ cells in the neural tube, a similar effect is likely occurring in the myogenic precursors of the heart. Lbx1 has been proposed to be involved in the expression of downstream genes involved in guidance cue interpretation and the maintenance of migratory potential in cells (Brohmann et al., 2000; Schmitteckert et al., 2011). Previous studies have identified Lbx1 expression in myogenic precursors and the myocardium of the developing heart. Severe defects in heart looping typically result in early embryonic lethality making it hard to detect/study the consequences later in development (Schafer et al., 2003). The current study’s findings align with previous Lbx1 KO studies, where mice with normal inflow/outflow and heart looping display ventricular hyperplasia/hypertrophy resulting in thickened ventricles(Schafer et al., 2003). The hyperplastic/hypertrophic ventricles are attributable to decreased *Fhl2* expression following altered *Lbx1* expression (Fig. SX).

This is different from skeletal muscle, where PAX3 is needed for Lbx1 expression in migrating myoblasts (Mennerich et al., 1998). In chicks, increased *lbx1* expression leads to increases myoD synthesis (Mennerich and Braun, 2001). MyoD is often used to mark terminal skeletal muscle differentiation. While in *Xenopus*, overexpression of *lbx1* lead to a lack of differentiated muscle, subsequent gain-of-function experiments co-injecting *myoD*, and *lbx1* mRNA transcripts restored myocyte numbers through the repressive actions of *lbx1* on *myoD* (Martin and Harland, 2006). Here, we found that an increase of *Lbx1* expression in the *Lbx1*^*Δ/Δ*^ had no apparent effect on the migration of myogenic cells, although it did lead to a decrease in *myoD* mRNA expression (Fig. S3).

Of the differentially expressed genes, there are multiple transcription factors. These included: *Hox* genes family who have multifaceted roles in neuronal specification and connectivity (Philippidou and Dasen, 2013), *Irx4*, which contributes to neural patterning (Gomez-Skarmeta and Modolell, 2002; Rodriguez-Seguel et al., 2009), *Lhx4*, which specifies the trajectory of motor axons in the neural tube (Sharma et al., 1998), *Prdm8* and *Prdm10* genes essential for the dorsoventral patterning (Zannino and Sagerstrom, 2015) and *WT1* necessary for the development of the locomotor circuitry (Haque et al., 2018). As we observed differences in total neuron markers and the altered/ectopic expression of neuronal population markers, we wanted to determine if there were any consequences on the trajectory of axons peripherally. Therefore, we chose to look at peripherin, often used to mark peripherally extending neurons, and observed a lack of long-range peripherally coursing axons in the *Lbx1*^*Δ/Δ*^ mouse. This suggests a deficit in the number of interneurons forming successful long-range connections and signaling appropriately through the spinal cord. While peripherin is not the only intermediate filament found in these axons, the small stature of the E18.5 embryos and the embryonic lethality suggests that neuronal connectivity and stimulation are not occurring during the prenatal period.

AIS is sometimes referred to as an asynchronous neuro-osseous growth, meaning disproportional growth occurs between the nervous and skeletal systems (Lam et al., 2011; Lao et al., 2011; Porter, 2001). The asynchronous neuro-osseous growth theory is based on findings also described as functional tethering of the spinal cord, whereby during periods of rapid growth, the spinal cord is unable to grow in length as rapidly as the vertebrae and therefore creates an axis around which the spine twists and bends to compensate (Chu et al., 2008; Porter, 2001). The alteration of the spine and vertebrae length suggests that LBX1 dysfunction can interfere with the correct development of the vertebral column; moreover, abnormal skeletal growth patterns are a hallmark of AIS (Yim et al., 2012). Short spinal cord and vertebral canal have been linked to scoliosis (Porter, 2001).

Altogether, our study increases our understanding Lbx1s function in embryonic development and provides a model for studying loss/gain of function, following the deletion of the EH1 domain. Our findings also indicate that the correlation between Lbx1 and AIS maybe due to Lbx1’s role in the correct development of the neuronal circuitry and that many other genes linked to scoliotic phenotypes also form part of the same genetic network.

## Acknowledgments

The authors wish to thank Pr. Carmen Birchmeier and Dr. Thomas Müller for the Lbx1 antibody. We would like to thank Edward Moody, Susie Szakats and Michael Meier for feedback on drafts of the manuscript.

## References

Assaiante, C., Mallau, S., Jouve, J.L., Bollini, G., Vaugoyeau, M., 2012. Do adolescent idiopathic scoliosis (AIS) neglect proprioceptive information in sensory integration of postural control? PLoS One 7, e40646.

Blecher, R., Krief, S., Galili, T., Biton, I.E., Stern, T., Assaraf, E., Levanon, D., Appel, E., Anekstein, Y., Agar, G., Groner, Y., Zelzer, E., 2017. The Proprioceptive System Masterminds Spinal Alignment: Insight into the Mechanism of Scoliosis. Dev Cell 42, 388–399 e383.

Brohmann, H., Jagla, K., Birchmeier, C., 2000. The role of Lbx1 in migration of muscle precursor cells. Development 127, 437–445.

Bulman, M.P., Kusumi, K., Frayling, T.M., McKeown, C., Garrett, C., Lander, E.S., Krumlauf, R., Hattersley, A.T., Ellard, S., Turnpenny, P.D., 2000. Mutations in the human delta homologue, DLL3, cause axial skeletal defects in spondylocostal dysostosis. Nat Genet 24, 438–441.

Catela, C., Bilbao-Cortes, D., Slonimsky, E., Kratsios, P., Rosenthal, N., Te Welscher, P., 2009. Multiple congenital malformations of Wolf-Hirschhorn syndrome are recapitulated in Fgfrl1 null mice. Dis Model Mech 2, 283–294.

Chen, X., Chen, J., Xu, D., Zhao, S., Song, H., Peng, Y., 2017. Effects of Osteoglycin (OGN) on treating senile osteoporosis by regulating MSCs. BMC Musculoskelet Disord 18, 423.

Cheng, L., Samad, O.A., Xu, Y., Mizuguchi, R., Luo, P., Shirasawa, S., Goulding, M., Ma, Q., 2005. Lbx1 and Tlx3 are opposing switches in determining GABAergic versus glutamatergic transmitter phenotypes. Nat Neurosci 8, 1510–1515.

Chu, W.C., Lam, W.M., Ng, B.K., Tze-Ping, L., Lee, K.M., Guo, X., Cheng, J.C., Burwell, R.G., Dangerfield, P.H., Jaspan, T., 2008. Relative shortening and functional tethering of spinal cord in adolescent scoliosis - Result of asynchronous neuro-osseous growth, summary of an electronic focus group debate of the IBSE. Scoliosis 3, 8.

Dehni, G., Liu, Y., Husain, J., Stifani, S., 1995. TLE expression correlates with mouse embryonic segmentation, neurogenesis, and epithelial determination. Mech Dev 53, 369–381.

Delile, J., Rayon, T., Melchionda, M., Edwards, A., Briscoe, J., Sagner, A., 2019. Single cell transcriptomics reveals spatial and temporal dynamics of gene expression in the developing mouse spinal cord. Development 146.

Fadzan, M., Bettany-Saltikov, J., 2017. Etiological Theories of Adolescent Idiopathic Scoliosis: Past and Present. Open Orthop J 11, 1466–1489.

Gomez-Skarmeta, J.L., Modolell, J., 2002. Iroquois genes: genomic organization and function in vertebrate neural development. Curr Opin Genet Dev 12, 403–408.

Grauers, A., Wang, J., Einarsdottir, E., Simony, A., Danielsson, A., Akesson, K., Ohlin, A., Halldin, K., Grabowski, P., Tenne, M., Laivuori, H., Dahlman, I., Andersen, M., Christensen, S.B., Karlsson, M.K., Jiao, H., Kere, J., Gerdhem, P., 2015. Candidate gene analysis and exome sequencing confirm LBX1 as a susceptibility gene for idiopathic scoliosis. Spine J 15, 2239–2246.

Grimes, D.T., Boswell, C.W., Morante, N.F., Henkelman, R.M., Burdine, R.D., Ciruna, B., 2016. Zebrafish models of idiopathic scoliosis link cerebrospinal fluid flow defects to spine curvature. Science 352, 1341–1344.

Gross, M.K., Dottori, M., Goulding, M., 2002. Lbx1 specifies somatosensory association interneurons in the dorsal spinal cord. Neuron 34, 535–549.

Guo, L., Yamashita, H., Kou, I., Takimoto, A., Meguro-Horike, M., Horike, S., Sakuma, T., Miura, S., Adachi, T., Yamamoto, T., Ikegawa, S., Hiraki, Y., Shukunami, C., 2016. Functional Investigation of a Non-coding Variant Associated with Adolescent Idiopathic Scoliosis in Zebrafish: Elevated Expression of the Ladybird Homeobox Gene Causes Body Axis Deformation. PLoS Genet 12, e1005802.

Haque, F., Rancic, V., Zhang, W., Clugston, R., Ballanyi, K., Gosgnach, S., 2018. WT1-Expressing Interneurons Regulate Left-Right Alternation during Mammalian Locomotor Activity. The Journal of neuroscience : the official journal of the Society for Neuroscience 38, 5666–5676.

Hernandez-Miranda, L.R., Ibrahim, D.M., Ruffault, P.L., Larrosa, M., Balueva, K., Muller, T., Weerd, W., Stolte-Dijkstra, I., Hostra, R.M.W., Brunet, J.F., Fortin, G., Mundlos, S., Birchmeier, C., 2018. Mutation in LBX1/Lbx1 precludes transcription factor cooperativity and causes congenital hypoventilation in humans and mice. Proc Natl Acad Sci U S A 115, 13021–13026.

Heywood, J.L., McEntee, G.M., Stickland, N.C., 2005. In ovo neuromuscular stimulation alters the skeletal muscle phenotype of the chick. J Muscle Res Cell Motil 26, 49–56.

Higuchi, M., Tomioka, M., Takano, J., Shirotani, K., Iwata, N., Masumoto, H., Maki, M., Itohara, S., Saido, T.C., 2005. Distinct mechanistic roles of calpain and caspase activation in neurodegeneration as revealed in mice overexpressing their specific inhibitors. J Biol Chem 280, 15229–15237.

Jen, J.C., Chan, W.M., Bosley, T.M., Wan, J., Carr, J.R., Rub, U., Shattuck, D., Salamon, G., Kudo, L.C., Ou, J., Lin, D.D., Salih, M.A., Kansu, T., Al Dhalaan, H., Al Zayed, Z., MacDonald, D.B., Stigsby, B., Plaitakis, A., Dretakis, E.K., Gottlob, I., Pieh, C., Traboulsi, E.I., Wang, Q., Wang, L., Andrews, C., Yamada, K., Demer, J.L., Karim, S., Alger, J.R., Geschwind, D.H., Deller, T., Sicotte, N.L., Nelson, S.F., Baloh, R.W., Engle, E.C., 2004. Mutations in a human ROBO gene disrupt hindbrain axon pathway crossing and morphogenesis. Science 304, 1509–1513.

Karner, C.M., Long, F., Solnica-Krezel, L., Monk, K.R., Gray, R.S., 2015. Gpr126/Adgrg6 deletion in cartilage models idiopathic scoliosis and pectus excavatum in mice. Hum Mol Genet 24, 4365–4373.

Koop, K.E., MacDonald, L.M., Lobe, C.G., 1996. Transcripts of Grg4, a murine groucho-related gene, are detected in adjacent tissues to other murine neurogenic gene homologues during embryonic development. Mech Dev 59, 73–87.

Kruger, M., Schafer, K., Braun, T., 2002. The homeobox containing gene Lbx1 is required for correct dorsal-ventral patterning of the neural tube. J Neurochem 82, 774–782.

Lam, T.P., Hung, V.W., Yeung, H.Y., Tse, Y.K., Chu, W.C., Ng, B.K., Lee, K.M., Qin, L., Cheng, J.C., 2011. Abnormal bone quality in adolescent idiopathic scoliosis: a case-control study on 635 subjects and 269 normal controls with bone densitometry and quantitative ultrasound. Spine (Phila Pa 1976) 36, 1211–1217.

Lao, L.F., Shen, J.X., Chen, Z.G., Wang, Y.P., Wen, X.S., Qiu, G.X., 2011. Uncoupled neuro-osseous growth in adolescent idiopathic scoliosis? A preliminary study of 90 adolescents with whole-spine three-dimensional magnetic resonance imaging. European spine journal : official publication of the European Spine Society, the European Spinal Deformity Society, and the European Section of the Cervical Spine Research Society 20, 1081–1086.

Leonard, D.G., Gorham, J.D., Cole, P., Greene, L.A., Ziff, E.B., 1988. A nerve growth factor-regulated messenger RNA encodes a new intermediate filament protein. J Cell Biol 106, 181–193.

Liu, S., Wu, N., Zuo, Y., Zhou, Y., Liu, J., Liu, Z., Chen, W., Liu, G., Chen, Y., Chen, J., Lin, M., Zhao, Y., Ming, Y., Yuan, T., Li, X., Xia, Z., Yang, X., Ma, Y., Zhang, J., Shen, J., Li, S., Wang, Y., Zhao, H., Yu, K., Zhao, Y., Weng, X., Qiu, G., Wu, Z., 2017. Genetic Polymorphism of LBX1 Is Associated With Adolescent Idiopathic Scoliosis in Northern Chinese Han Population. Spine (Phila Pa 1976) 42, 1125–1129.

Londono, D., Kou, I., Johnson, T.A., Sharma, S., Ogura, Y., Tsunoda, T., Takahashi, A., Matsumoto, M., Herring, J.A., Lam, T.P., Wang, X., Tam, E.M., Song, Y.Q., Fan, Y.H., Chan, D., Cheah, K.S., Qiu, X., Jiang, H., Huang, D., Japanese Scoliosis Clinical Research, G., Group, T.I.C., International Consortium for Scoliosis, G., Su, P., Sham, P., Cheung, K.M., Luk, K.D., Gordon, D., Qiu, Y., Cheng, J., Tang, N., Ikegawa, S., Wise, C.A., 2014. A meta-analysis identifies adolescent idiopathic scoliosis association with LBX1 locus in multiple ethnic groups. J Med Genet 51, 401–406.

Loomes, K.M., Stevens, S.A., O’Brien, M.L., Gonzalez, D.M., Ryan, M.J., Segalov, M., Dormans, N.J., Mimoto, M.S., Gibson, J.D., Sewell, W., Schaffer, A.A., Nah, H.D., Rappaport, E.F., Pratt, S.C., Dunwoodie, S.L., Kusumi, K., 2007. Dll3 and Notch1 genetic interactions model axial segmental and craniofacial malformations of human birth defects. Developmental dynamics : an official publication of the American Association of Anatomists 236, 2943–2951.

Lu, D.C., Niu, T., Alaynick, W.A., 2015. Molecular and cellular development of spinal cord locomotor circuitry. Front Mol Neurosci 8, 25.

Martin, B.L., Harland, R.M., 2006. A novel role for lbx1 in Xenopus hypaxial myogenesis. Development 133, 195–208.

Masselink, W., Masaki, M., Sieiro, D., Marcelle, C., Currie, P.D., 2017. Phosphorylation of Lbx1 controls lateral myoblast migration into the limb. Dev Biol 430, 302–309.

Mennerich, D., Braun, T., 2001. Activation of myogenesis by the homeobox gene Lbx1 requires cell proliferation. EMBO J 20, 7174–7183.

Mennerich, D., Schafer, K., Braun, T., 1998. Pax-3 is necessary but not sufficient for lbx1 expression in myogenic precursor cells of the limb. Mech Dev 73, 147–158.

Mizuhara, E., Nakatani, T., Minaki, Y., Sakamoto, Y., Ono, Y., 2005. Corl1, a novel neuronal lineage-specific transcriptional corepressor for the homeodomain transcription factor Lbx1. J Biol Chem 280, 3645–3655.

Muhr, J., Andersson, E., Persson, M., Jessell, T.M., Ericson, J., 2001. Groucho-mediated transcriptional repression establishes progenitor cell pattern and neuronal fate in the ventral neural tube. Cell 104, 861–873.

Muller, T., Brohmann, H., Pierani, A., Heppenstall, P.A., Lewin, G.R., Jessell, T.M., Birchmeier, C., 2002. The homeodomain factor lbx1 distinguishes two major programs of neuronal differentiation in the dorsal spinal cord. Neuron 34, 551–562.

Pagliardini, S., Ren, J., Gray, P.A., Vandunk, C., Gross, M., Goulding, M., Greer, J.J., 2008. Central respiratory rhythmogenesis is abnormal in lbx1-deficient mice. The Journal of neuroscience : the official journal of the Society for Neuroscience 28, 11030–11041.

Pfaff, S.L., Mendelsohn, M., Stewart, C.L., Edlund, T., Jessell, T.M., 1996. Requirement for LIM homeobox gene Isl1 in motor neuron generation reveals a motor neuron-dependent step in interneuron differentiation. Cell 84, 309–320.

Philippidou, P., Dasen, J.S., 2013. Hox genes: choreographers in neural development, architects of circuit organization. Neuron 80, 12–34.

Porter, R.W., 2001. The pathogenesis of idiopathic scoliosis: uncoupled neuro-osseous growth? European spine journal : official publication of the European Spine Society, the European Spinal Deformity Society, and the European Section of the Cervical Spine Research Society 10, 473–481.

Raudvere, U., Kolberg, L., Kuzmin, I., Arak, T., Adler, P., Peterson, H., Vilo, J., 2019. g:Profiler: a web server for functional enrichment analysis and conversions of gene lists (2019 update). Nucleic Acids Res 47, W191–W198.

Richardson, L., Venkataraman, S., Stevenson, P., Yang, Y., Moss, J., Graham, L., Burton, N., Hill, B., Rao, J., Baldock, R.A., Armit, C., 2014. EMAGE mouse embryo spatial gene expression database: 2014 update. Nucleic Acids Res 42, D835–844.

Rigueur, D., Lyons, K.M., 2014. Whole-mount skeletal staining. Methods Mol Biol 1130, 113–121.

Robinson, M.D., McCarthy, D.J., Smyth, G.K., 2010. edgeR: a Bioconductor package for differential expression analysis of digital gene expression data. Bioinformatics 26, 139-140.

Rodriguez-Seguel, E., Alarcon, P., Gomez-Skarmeta, J.L., 2009. The Xenopus Irx genes are essential for neural patterning and define the border between prethalamus and thalamus through mutual antagonism with the anterior repressors Fezf and Arx. Dev Biol 329, 258–268.

Sammeta, N., Hardin, D.L., McClintock, T.S., 2010. Uncx regulates proliferation of neural progenitor cells and neuronal survival in the olfactory epithelium. Mol Cell Neurosci 45, 398–407.

Schafer, K., Neuhaus, P., Kruse, J., Braun, T., 2003. The homeobox gene Lbx1 specifies a subpopulation of cardiac neural crest necessary for normal heart development. Circ Res 92, 73–80.

Schmitteckert, S., Ziegler, C., Kartes, L., Rolletschek, A., 2011. Transcription factor lbx1 expression in mouse embryonic stem cell-derived phenotypes. Stem Cells Int 2011, 130970.

Schnerwitzki, D., Perry, S., Ivanova, A., Caixeta, F.V., Cramer, P., Gunther, S., Weber, K., Tafreshiha, A., Becker, L., Vargas Panesso, I.L., Klopstock, T., Hrabe de Angelis, M., Schmidt, M., Kullander, K., Englert, C., 2018. Neuron-specific inactivation of Wt1 alters locomotion in mice and changes interneuron composition in the spinal cord. Life Sci Alliance 1, e201800106.

Sharma, K., Sheng, H.Z., Lettieri, K., Li, H., Karavanov, A., Potter, S., Westphal, H., Pfaff, S.L., 1998. LIM homeodomain factors Lhx3 and Lhx4 assign subtype identities for motor neurons. Cell 95, 817–828.

Sieber, M.A., Storm, R., Martinez-de-la-Torre, M., Muller, T., Wende, H., Reuter, K., Vasyutina, E., Birchmeier, C., 2007. Lbx1 acts as a selector gene in the fate determination of somatosensory and viscerosensory relay neurons in the hindbrain. The Journal of neuroscience : the official journal of the Society for Neuroscience 27, 4902–4909.

Singh, M.K., Petry, M., Haenig, B., Lescher, B., Leitges, M., Kispert, A., 2005. The T-box transcription factor Tbx15 is required for skeletal development. Mech Dev 122, 131–144.

Takahashi, Y., Kou, I., Takahashi, A., Johnson, T.A., Kono, K., Kawakami, N., Uno, K., Ito, M., Minami, S., Yanagida, H., Taneichi, H., Tsuji, T., Suzuki, T., Sudo, H., Kotani, T., Watanabe, K., Chiba, K., Hosono, N., Kamatani, N., Tsunoda, T., Toyama, Y., Kubo, M., Matsumoto, M., Ikegawa, S., 2011. A genome-wide association study identifies common variants near LBX1 associated with adolescent idiopathic scoliosis. Nat Genet 43, 1237–1240.

Takeda, K., Kou, I., Mizumoto, S., Yamada, S., Kawakami, N., Nakajima, M., Otomo, N., Ogura, Y., Miyake, N., Matsumoto, N., Kotani, T., Sudo, H., Yonezawa, I., Uno, K., Taneichi, H., Watanabe, K., Shigematsu, H., Sugawara, R., Taniguchi, Y., Minami, S., Nakamura, M., Matsumoto, M., Japan Early Onset Scoliosis Research, G., Watanabe, K., Ikegawa, S., 2018. Screening of known disease genes in congenital scoliosis. Mol Genet Genomic Med 6, 966–974.

Tonami, K., Hata, S., Ojima, K., Ono, Y., Kurihara, Y., Amano, T., Sato, T., Kawamura, Y., Kurihara, H., Sorimachi, H., 2013. Calpain-6 deficiency promotes skeletal muscle development and regeneration. PLoS Genet 9, e1003668.

Troy, C.M., Greene, L.A., Shelanski, M.L., 1992. Neurite outgrowth in peripherin-depleted PC12 cells. J Cell Biol 117, 1085–1092.

Volk, A.E., Carter, O., Fricke, J., Herkenrath, P., Poggenborg, J., Borck, G., Demant, A.W., Ivo, R., Eysel, P., Kubisch, C., Neugebauer, A., 2011. Horizontal gaze palsy with progressive scoliosis: three novel ROBO3 mutations and descriptions of the phenotypes of four patients. Mol Vis 17, 1978–1986.

Warnes, G.R., Bolker, B., Bonebakker, L., Gentleman, R., Huber, W., Liaw, A., Lumley, T., Maechler, M., Magnusson, A., Moeller, S., 2009. gplots: Various R programming tools for plotting data. R package version 2, 1.

Weinstein, S.L., Dolan, L.A., Cheng, J.C., Danielsson, A., Morcuende, J.A., 2008. Adolescent idiopathic scoliosis. Lancet 371, 1527–1537.

Xu, J.F., Yang, G.H., Pan, X.H., Zhang, S.J., Zhao, C., Qiu, B.S., Gu, H.F., Hong, J.F., Cao, L., Chen, Y., Xia, B., Bi, Q., Wang, Y.P., 2015. Association of GPR126 gene polymorphism with adolescent idiopathic scoliosis in Chinese populations. Genomics 105, 101–107.

Yagnik, G., Ghuman, A., Kim, S., Stevens, C.G., Kimonis, V., Stoler, J., Sanchez-Lara, P.A., Bernstein, J.A., Naydenov, C., Drissi, H., Cunningham, M.L., Kim, J., Boyadjiev, S.A., 2012. ALX4 gain-of-function mutations in nonsyndromic craniosynostosis. Hum Mutat 33, 1626–1629.

Yim, A.P., Yeung, H.Y., Hung, V.W., Lee, K.M., Lam, T.P., Ng, B.K., Qiu, Y., Cheng, J.C., 2012. Abnormal skeletal growth patterns in adolescent idiopathic scoliosis--a longitudinal study until skeletal maturity. Spine (Phila Pa 1976) 37, E1148–1154.

Zannino, D.A., Sagerstrom, C.G., 2015. An emerging role for prdm family genes in dorsoventral patterning of the vertebrate nervous system. Neural Dev 10, 24.

