## Supplementary data for "Characterization of a novel *Lbx1* mouse loss of function strain"

Megan J. Wilson*

Department of Anatomy, Otago School of Medical Sciences, University of Otago, 9054

Dunedin, New Zealand

*Corresponding author:

**Supplementary Files:**

**File 1: List of differentially expressed genes.**

**File 2: gProfiler terms and gene associations.**

**Supplementary Tables**

**Supplementary Table 1. List of primers used for PCR.** Primers sorted by alphabetical order.

| ID primer | Purpose | Fwd sequence | Rv sequence |
|---|---|---|---|
| Actin | RT-qPCR | GGCTGTATTCCCCTCCATCG | CCAGTTGGTAACAATGCCATGT |
| Brn3a | RT-qPCR | GAAATGCCTGCTTTGTTGTGTA | CCATGCATGATTTCAGTCTTGT |
| Clic6 | RT-qPCR | GAGGTACCCTAAGCTGGGAACT | ATTTGCATCCTTCTTGGTGTTT |
| Gapdh | RT-qPCR | AAGGTCATCCCAGAGCTGAA | CTGCTTCACCACCTTCTTGA |
| Ide | RT-qPCR | ACTAACCTGGTGGTGAAGCTGT | AAAGTTGCCTGAGATGCTCTTC |
| Lbx1 | ISH | CTTCACCAACCACCAGATCTAC | CCACGTCGATCTCTTCATCTTC |
| | PCR genotyping | ACGAGGCCGAGATGACTTC | CCCACACAGCGAGTAACTT |
| | RT-qPCR | GCGATGGGATGACCATCTTT | GCGTTTCTCCAACTCGTAGAT |
| | Sequencing | ATGACTTCCAAGGAGGACG | / |
| Lhx2 | RT-qPCR | AACGTGTAACAAGATGCTGACG | CATGGTTAAAGTGTGCTGGGTA |
| Lhx9 | ISH | Gift from Pr. Peter Koopman and Dr. Dagmar Wilhelm, Genes Dev. 2002 Jul 15; 16(14): 1839–1851 | |
| | RT-qPCR | GCCAAGGACGGTAGCATTTA | AGCTCAGATGGTAGACAGAGT |
| Lmx1b | ISH | CTGTGCAAGGGTGACTATGAGA | ATATCGTGGAAGATGGAGTCGT |
| | RT-qPCR | CAGCAGCGAAGAGCTTTCAA | GTCTCTCGGACCTTCCGACA |
| Lrrc17 | RT-qPCR | CTCTGTGCCACAATTACGTGTT | TTTTGTTGGATGACAGGTCAAG |
| Ogn | RT-qPCR | TAAAAAGCTGACTGCCAAGGA | AAGCTTTGGAGGAAGAACTGG |
| Pax2 | ISH | TAATGGAGACTCCCAGAGTGGT | GTAACTAGTGGCGGTCATAGGC |
| | RT-qPCR | TGGCTGTGTCAGCAAAATCCT | ATTCGGCAATCTTGTCCACCA |
| Pax3 | ISH | GGCTCCGATATTGACTCTGAAC | TTCGAAGGAATAGTGCTTTGGT |
| | RT-qPCR | TCAGAGACAAATTGCTCAAGGA | AGCCTTTTTCTCGCTTTCTTCT |
| Prdm8 | RT-qPCR | AATGGTTCCTCAGAAGGTCTCA | AGTCAGTTCTTTCCCGTACCAA |
| Prrx1 | RT-qPCR | AATGTACGCATGTTCTGGTCAC | TTATTGTTTGGGTTGGGTAAGG |
| Robo3 | RT-qPCR | TCATTCCAATGTGAGACCAAAG | CATGGGTTGGAGTGACTGACTA |
| Six2 | RT-qPCR | GGTATCCTTTGGGGATTCATTT | GGGTGGTGGTGGCTTATAATTT |
| Slc15a2 | RT-qPCR | GCTAAATCCCTTTCTGGTCCTT | ATCATACCAACAGCCATTTTCC |
| Tbx15 | RT-qPCR | CATTCAGTGTGCCATCTTTGAT | GTCTTGCTGTCCACTCTCTCCT |
| Uncx | RT-qPCR | GCGTTCAATGAGAGCCACTAC | GATTTTGGAACCAGACCTGAAC |
| Wt1 | RT-qPCR | GGAGCTACCTTAAAGGGAATGG | CGTGTGGTTCTCACTCTCATA |

**Supplementary Table 2. Align summary of TopHat mapping.** Statistics presenting the number of reads that mapped to the genome (mm9).

| | Left reads | | Right reads | | |
|---|---|---|---|---|---|
| **ID** | **Input** | **Mapped** | **Input** | **Mapped** | **Aligned pairs** |
| **Control - 1** | 24,037,892 | 22,792,206 (94.8% of input) | 24,037,892 | 21,226,401 (88.3% of input) | 20,646,450 (85.6% concordant pair alignment rate) |
| **Control - 2** | 20,222,001 | 19,146,209 (94.7% of input) | 20,222,001 | 17,883,701 (88.4% of input) | 17,360,718 (85.6% concordant pair alignment rate) |
| **Lbx1Δ - 1** | 20,276,958 | 19,280,476 (95.1% of input) | 20,276,958 | 17,617347 (86.9% of input) | 17141525 (84.2% concordant pair alignment rate) |
| **Lbx1Δ - 2** | 20,106,498 | 19,098,089 (95.0% of input) | 20,106,498 | 17,748,677 (88.3% of input) | 17,257,469 (85.6% concordant pair alignment rate) |

**Supplementary Table 3. Gene list of the RNA seq results.** List of the differentially expressed genes in Lbx1 $^{\Delta/\Delta}$ E12.5 neural tubes compared to wild types with a fold change (logFC) either below -1 or above 1 (grey**)**. Adj.P.Val: adjusted p-value.

| refSeq | logFC | P.Value | adj.P.Val | gene ID |
|---|---|---|---|---|
| NM_144783 | -2.10 | 1.34E-12 | 1.68E-09 | *Wt1* |
| NM_010055 | -1.67 | 5.53E-07 | 9.80E-05 | *Dlx3* |
| NM_172469 | -1.64 | 2.99E-09 | 1.24E-06 | *Clic6* |
| NM_172925 | -1.60 | 1.31E-06 | 1.95E-04 | *Klhl31* |
| NM_021301 | -1.56 | 9.95E-13 | 1.42E-09 | *Slc15a2* |
| NM_178924 | -1.54 | 6.48E-05 | 4.43E-03 | *Upk1b* |
| NM_001081281 | -1.52 | 3.94E-06 | 4.85E-04 | *Trim55* |
| NM_001033621 | -1.50 | 2.93E-05 | 2.44E-03 | *Myot* |
| NM_010769 | -1.49 | 6.10E-05 | 4.26E-03 | *Matn1* |
| NM_001081157 | -1.38 | 3.18E-07 | 6.11E-05 | *Lmod3* |
| NM_022322 | -1.25 | 7.78E-05 | 5.04E-03 | *Tnmd* |
| NM_026841 | -1.24 | 2.38E-04 | 1.11E-02 | *Prr32* |
| NM_010258 | -1.21 | 6.12E-04 | 2.24E-02 | *Gata6* |
| NM_054049 | -1.18 | 6.16E-04 | 2.25E-02 | *Osr2* |
| NM_007442 | -1.16 | 2.32E-13 | 4.29E-10 | *Alx4* |
| NM_008312 | -1.16 | 1.41E-03 | 3.86E-02 | *Htr2c* |
| NM_011380 | -1.15 | 9.84E-10 | 5.46E-07 | *Six2* |
| NM_013738 | -1.14 | 1.22E-03 | 3.59E-02 | *Plek2* |
| NM_007733 | -1.11 | 8.23E-06 | 8.56E-04 | *Col19a1* |
| NM_001166067 | -1.11 | 5.46E-04 | 2.10E-02 | *Slc4a5* |
| NM_009710 | -1.10 | 1.29E-03 | 3.68E-02 | *Art1* |
| NM_008760 | -1.10 | 2.58E-13 | 4.29E-10 | *Ogn* |
| NM_175093 | -1.09 | 6.89E-04 | 2.40E-02 | *Trib3* |
| NM_007504 | -1.07 | 1.15E-07 | 2.73E-05 | *Atp2a1* |
| NM_013930 | -1.07 | 5.43E-04 | 2.10E-02 | *Aass* |
| NM_030728 | -1.03 | 2.22E-04 | 1.08E-02 | *Cemip* |
| NM_001081087 | -1.02 | 4.22E-09 | 1.68E-06 | *Klhl41* |
| NM_007389 | -1.01 | 4.80E-05 | 3.58E-03 | *Chrna1* |
| NM_009323 | -1.01 | 1.12E-13 | 2.79E-10 | *Tbx15* |
| NM_010691 | 1.87 | 1.13E-52 | 1.13E-48 | *Lbx1* |
| NM_009654 | 1.63 | 3.00E-07 | 5.88E-05 | *Alb* |
| NM_028862 | 1.50 | 2.24E-05 | 1.93E-03 | *Rnf145* |
| NR_037698 | 1.37 | 6.08E-04 | 2.24E-02 | *1500009C09Rik* |
| NM_001136058 | 1.20 | 1.78E-07 | 3.95E-05 | *Crmp1* |
| NM_173410 | 1.19 | 1.27E-03 | 3.67E-02 | *Gpr26* |

| | | | | |
|---|---|---|---|---|
| NM_001080819 | 1.16 | 2.97E-04 | 1.31E-02 | *Arid1a* |
| NM_013625 | 1.16 | 1.39E-10 | 9.92E-08 | *Pafah1b1* |
| NM_001355181 | 1.06 | 1.80E-04 | 9.27E-03 | *Rnf38* |
| NM_001163565 | 1.06 | 3.27E-05 | 2.68E-03 | *Ptpn5* |
| NM_001009930 | 1.03 | 7.58E-04 | 2.56E-02 | *Brsk2* |
| NM_011436 | 1.00 | 1.86E-03 | 4.64E-02 | *Sorl1* |

**Supplementary Table 4. Differentially expressed genes in RNA-seq involved in neuronal identity.**

| gene ID | logFC | adj.p.value |
| --- | --- | --- |
| *Hoxa3* | 0.32 | 4.34E-02 |
| *Hoxa6* | 0.74 | 1.53E-05 |
| *Hoxb5* | 0.33 | 1.08E-02 |
| *Hoxb6* | 0.42 | 4.95E-04 |
| *Hoxb8* | 0.27 | 3.86E-02 |
| *Hoxb9* | 0.34 | 3.68E-03 |
| *Hoxd3* | 0.37 | 3.82E-03 |
| *Hoxd4* | 0.50 | 2.94E-04 |
| *Irx4* | -0.88 | 1.75E-02 |
| *Lhx4* | 0.64 | 3.15E-02 |
| *Prdm10* | 0.57 | 1.14E-02 |
| *Prdm8* | 0.77 | 9.80E-05 |
| *WT1* | -2.10 | 1.68E-09 |

**Supplementary Table 5: Genes with links to human scoliosis phenotypes that are differentially expressed in the *Lbx1*$^{\Delta/\Delta}$ neural tube.**

| Gene name | $Log_2FC$ in *Lbx1*$^{\Delta/\Delta}$ | Adolescent idiopathic scoliosis | Congenital Scoliosis | Scoliosis phenotype, co-morbidity |
|---|---|---|---|---|
| Neuronal | | | | |
| *ROBO3* | 0.57 | | | Horizontal gaze palsy with progressive scoliosis (Jen et al., 2004) |
| Notch signalling | | | | |
| *NOTCH1* | 0.38 | | (Sparrow et al., 2012) | |
| *LFNG* | 0.37 | | (Takeda et al., 2018) | |
| *DLL3* | 0.34 | | | Kyphoscoliosis (Bulman et al., 2000; Giampietro et al., 2006) |
| *MEOX1* | -0.55 | ?notch? | | Klippel-Feil Syndrome 2 {Bayrakli, 2013 #498} |
| *SALL1* | 0.34 | | | Facio-auriculo-vertebral spectrum (Keegan et al., 2001) |
| *HES5* | 0.53 | | | |
| Extracellular matrix proteins | | | | |
| *COL3A1, COL5A2, COL9A3, COL11A1, COLL12A1* | -0.31, -0.46, -0.31, -0.47, -0.46 | (Haller et al., 2016) | | |
| *FBN1, FBN2* | -0.42, -0.43 | (Buchan et al., 2014; Sheng et al., 2019) | (Lin et al., 2020) | |
| *MATN1* | | (Zhang et al., 2014) | | |
| *MYOT* | -1.50 | | | Myofibrillar myopathy (Schessl et al., 2014; Selcen and Engel, 2004) |
| Transcription factors | | | | |
| *HOXB8* | | (Fendri et al., 2013) | | |
| *BMPR1a* | -0.30 | | | 10q22p23 deletion syndrome (Breckpot et al., 2012) |
| *CBL* | 0.39 | | | Noonan syndrome (Tartaglia et al., 2011) |
| *ARID1a* | 1.15 | | | Coffin-siris syndrome (Kosho et al., 2014) |
| *LBX1* | 1.88 | (Gao et al., 2013; Londono et al., 2014; Man et al., 2019; Nada et al., 2018; Takahashi et al., 2011; Zhu et al., 2015) | | |

5'-3' WT *Lbx1* sequence
gaggccgag**atg**acttccaaggaggacggcaaggcggcgccaggggaggagcggcgacgcagccctctggaccacctgccgccgcccgccaactc
caac**aagccgc**tgacgccgttcagcatcgaggacatcctcaacaagccgtccgtgcggagaagttactcgctgtgtggggcggcgcacctgctgg
cggccgcggacaagcacgcgccgggcggcttgcccctggcgggccgcgctctgctctcgcagacctcgcctctctgcgccttggaggagctcgcc
agcaagacctttaaggggctggaggtcagcgtcctgcaggcagccgaaggccgcgatgggatgaccatctttgggcagaggcagacgcccaagaa
acggcgaaaatcacgcacggccttcaccaaccaccagatctacgagttggagaaacgctttctataccagaagtacctgtccccggcagatcgcg
accaaattgcgcagcagctgggcctcaccaatgcacaggtcatcacctggttccagaaccggcgcgccaagctcaagcgggacctagaggagatg
aaggccgacgtggagtctgccaagaaactgggccccagcgggcagatggacatcgtggcgctggccgaactcgagcagaactcggaggcttcggg
cggtggcggcggcggtggctgcggcagggctaagtctaggccgggttctcctgcgctgcccccaggcgccccgcaggccccgggcggaggaccct
tgcagctctcgcccgcctctccactcacggaccagcgggccagcagccaggactgctcagaggatgaggaagatgaagagatcgacgtggacgat
tgagctgcg

5'3' WT LBX1 protein
Met T S K E D G K A A P G E E R R R S P L D H L P P P A N S N K P L T P F S I E D I L N K P S V
R R S Y S L C G A A H L L A A A D K H A P G G L P L A G R A L L S Q T S P L C A L E E L A S K T
F K G L E V S V L Q A A E G R D G Met T I F G Q R Q T P K K R R K S R T A F T N H Q I Y E L E K
R F L Y Q K Y L S P A D R D Q I A Q Q L G L T N A Q V I T W F Q N R R A K L K R D L E E Met K A
D V E S A K K L G P S G Q Met D I V A L A E L E Q N S E A S G G G G G G G C G R A K S R P G S P
A L P P G A P Q A P G G G P L Q L S P A S P L T D Q R A S S Q D C S E D E E D E E I D V D D
Stop

5'-3' *Lbx1*△
gaggccgag**atg**acttccaaggaggacggcaaggcggcgccaggggaggagcggcgacgcagccctctggaccacctgccgccgcccgccaactcc
aac**tga**cgccgttcagcatcgaggacatcctcaacaagccgtccgtgcggagaagttactcgctgtgtggggcggcgcacctgctggcggccgcgg
acaagcacgcgccgggcggcttgcccctggcgggccgcgctctgctctcgcagacctcgcctctctgcgccttggaggagctcgccagcaagacct
ttaaggggctggaggtcagcgtcctgcaggcagccgaaggccgcg**atg**ggatgaccatctttgggcagaggcagacgcccaagaaacggcgaaaat
cacgcacggccttcaccaaccaccagatctacgagttggagaaacgctttctataccagaagtacctgtccccggcagatcgcgaccaaattgcgc
agcagctgggcctcaccaatgcacaggtcatcacctggttccagaaccggcgcgccaagctcaagcgggacctagaggagatgaaggccgacgtgg
agtctgccaagaaactgggccccagcgggcagatggacatcgtggcgctggccgaactcgagcagaactcggaggcttcgggcggtggcggcggcg
gtggctgcggcagggctaagtctaggccgggttctcctgcgctgcccccaggcgccccgcaggccccgggcggaggacccttgcagctctcgcccg
cctctccactcacggaccagcgggccagcagccaggactgctcagaggatgaggaagatgaagagatcgacgtggacgattgagctgcg

5'-3' LBX1△
Met T I F G Q R Q T P K K R R K S R T A F T N H Q I Y E L E K R F L Y Q K Y L S P A D R D Q I A
Q Q L G L T N A Q V I T W F Q N R R A K L K R D L E E Met K A D V E S A K K L G P S G Q Met D I
V A L A E L E Q N S E A S G G G G G G G C G R A K S R P G S P A L P P G A P Q A P G G G P L Q L
S P A S P L T D Q R A S S Q D C S E D E E D E E I D V D D Stop

5'-3' LBX1
Met T S K E D G K A A P G E E R R R S P L D H L P P P A N S N **Stop**

**Figure S1 *Lbx1* mRNA and protein sequences.** A. The WT transcript sequence has the ATG highlighted in red for the first methionine. In green is the 7 bp sequence deleted by CRISPR-CAS9 editing.

B. *Lbx1Δ* predicted mRNA transcript. Highlighted in orange is a new stop codon created by the genome edit. This is predicted to results in two protein products. A short protein and a longer truncated LBX1 protein, missing the N-terminal end of the protein. Translation product from the alternative ATG (blue) is shown.

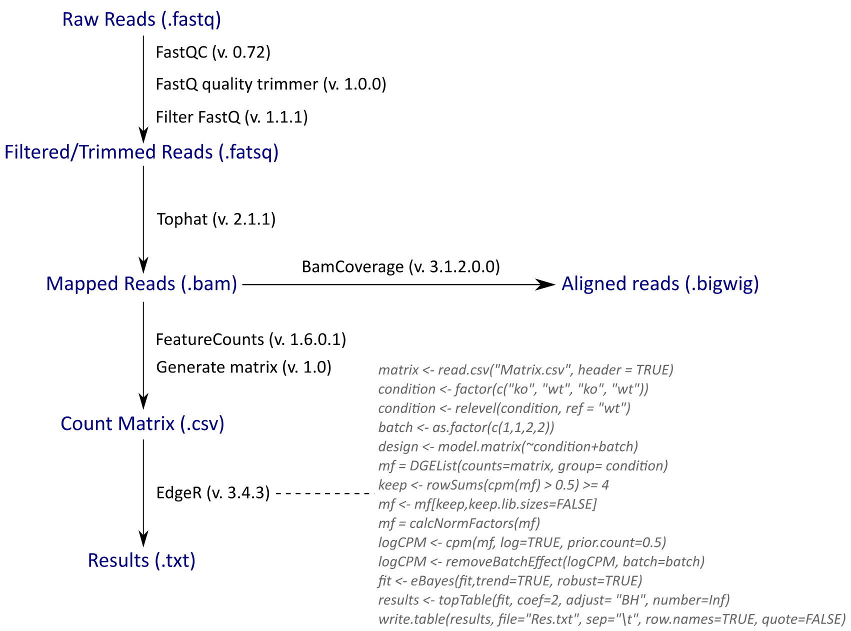

**Figure S2. RNA-sequencing pipeline.** Sequenced data were filtered and trimmed with FastQ quality trimmer and filter FastQ. The reads were then mapped to the reference genome mm9 (UCSC) using Tophat. The mapped reads were then either aligned to the genome using BamCoverage or processed using FeatureCounts and Generate matrix. For the differential gene expression analysis, edgeR was used, and the batch effect removed. Only significantly differentially expressed gene with an adjusted p-value <0.05, obtained with the FDR Benjamini Hochberg method, were taken for further analysis.

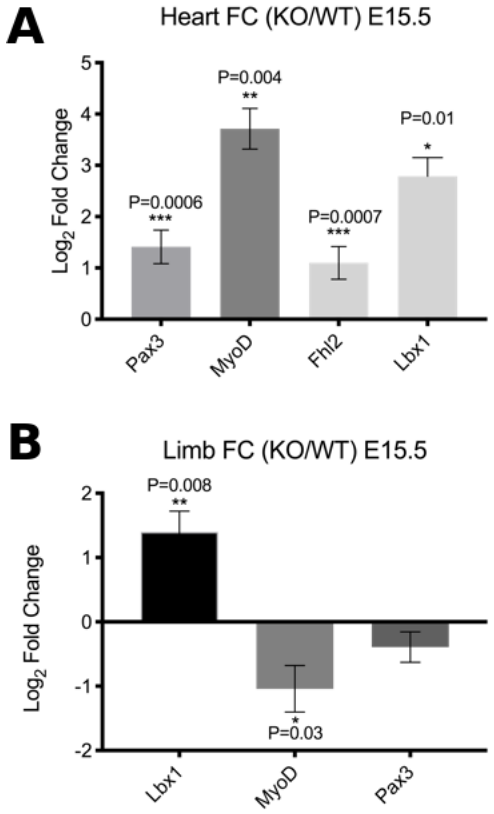

**Figure S3. Quantitative PCR analysis for heart and limb tissues.** Fold change mRNA levels between WT and *Lbx1*$^{\Delta/\Delta}$ limb and heart tissues at E15.5 (n = 5-7). Data is shown as mean +/- SEM, one-sample t-test with Wilcoxon test P < 0.05 (*), P < 0.001 (**), P < 0.0001 (***).

| dll | dI2 | dI3 | dI4 | dI5 | dI6 | v0 | v1 | v2a | v2b | MN | v3 | |
|---|---|---|---|---|---|---|---|---|---|---|---|---|
| | | | Lbx1 | Lbx1 | Lbx1 | | | | | | | UP |
| | | | | | Wt1 | | | | | | | DOWN |
| | Neurog2 | Neurog2 | Neurog2 | Neurog2 | | | Neurog2 | | Neurog2 | Neurog2 | | |
| | | | Neurod2 | | | Neurod2 | Neurod2 | Neurod2 | | | Neurod2 | |
| | | Sv2c | sv2c | | | | Sv2c | | | | Sv2c | |
| | HMX2 | | | | | | | HMX2 | | | | |
| | HMX3 | | | | | | | HMX3 | | | | |
| Uncx | Uncx | Uncx | Uncx | | | Uncx | | | | | | |
| | | | | Hoxb6 | | Hoxb9 | | | | | | |
| | | | | | | | | Igfbp2 | | | Igfbpl1 | |
| | | | | | | | | Hes5 | | Hes5 | | |
| | | | | | | | | Lhx4 | | Lhx4 | | |
| | | Pou3f2 | Pou3f2 | | | | | | | | | |
| | | Zfhx2 | | | | Zfhx2 | | | Zfhx2 | Zfhx2 | | |
| | | | | | | | | | Dll3 | Dll3 | | |
| Barhl2 | | | | | | | Spock2 | | | | | |
| | | Otp | | | | | | | | | | |
| | | | Fstl1 | | | | | | | | | |
| | | | Chmp2b | | | | | | | | | |
| | | | Npy | | | | | | | | | |
| | | | Igsf8 | | | | | | | | | |
| | | | Klhl35 | | | | | | | | | |
| | | | Nhlh1 | | | | | | | | | |
| | | | Rnd2 | | | | | | | | | |
| | | | | Shank1 | | Robo3 | | | | | | |
| | | | | Fam57b | | Cntn2 | | | | | | |
| | | | | Notch1 | | | | | | | | |

**Figure S4. DEGs with increased mRNA expression (green), reduced expression (red) plotted by neuron type.** Gene data was obtained from a published knowledge matrix, where a score of 1 indicates that this gene contributes to the cell type (column labels dl1-v3) {Delile, 2019 #521}.
